# Polymer-tethered quenched fluorescent probes for enhanced imaging of tumor associated proteases

**DOI:** 10.1101/2024.05.06.592849

**Authors:** Martin Hadzima, Franco Faucher, Kristýna Blažková, Joshua J. Yim, Matteo Guerra, Shiyu Chen, Emily C. Woods, Ki Wan Park, Pavel Šácha, Vladimír Šubr, Libor Kostka, Tomáš Etrych, Pavel Majer, Jan Konvalinka, Matthew Bogyo

## Abstract

Fluorescence-based contrast agents enable real-time detection of solid tumors and their neovasculature, making them ideal for use in image-guided surgery. Several agents have entered late-stage clinical trials or secured FDA approval, suggesting they are likely to become standard of care in cancer surgeries. One of the key parameters to optimize in contrast agent is molecular size, which dictates much of the pharmacokinetic and pharmacodynamic properties of the agent. Here, we describe the development of a class of protease-activated quenched fluorescent probes in which a N-(2-hydroxypropyl)methacrylamide copolymer is used as the primary scaffold. This copolymer core provides a high degree of probe modularity to generate structures that cannot be achieved with small molecules and peptide probes. We used a previously validated cathepsin substrate and evaluated the effects of length and type of linker as well as positioning of the fluorophore/quencher pair on the polymer core. We found that the polymeric probes could be optimized to achieve increased over-all signal and tumor-to-background ratios compared to the reference small molecule probe. Our results also revealed multiple structure-activity relationship trends that can be used to design and optimize future optical imaging probes. Furthermore, they confirm that a hydrophilic polymer is an ideal scaffold for use in optical imaging contrast probes, allowing a highly modular design that enables efficient optimization to maximize probe accumulation and overall biodistribution properties.

---

Precise cancer resection during surgical procedures re-mains a challenge due to difficulties with detection of tumor margins in real-time with high specificity and accuracy. Fluorescent probes have emerged as powerful tools with the potential to revolutionize cancer treatment as they pro-vide contrast without the need for exposure to radiation.1–5 Furthermore, the U.S. Food and Drug Administration (FDA) approval of multiple fluorescence imaging systems for use during surgery has created an opportunity to rapidly implement new optical contrast agents into existing surgical workflows. While FDA-approved fluorescent agents such as indocyanine green5,6 and methylene blue5,7 can be used in high dose to visualize some types of solid tumors through passive uptake or exclusion of the free dyes, their imaging performance is limited by the absence of a specific targeting mechanism. Recently, the FDA approved the first affinity-based fluorescent probe for cancer imaging, OTL38 (CYTALUX).8,9 OTL38 is a folic acid derivative bearing a near-infrared (NIR) fluorophore that exhibits high affinity for folate receptor α (FRα) which is often overexpressed in multiple types of solid tumors. In addition, activity-based probes, relying on specific enzymatic activation within the target tissue, are now reaching late-stage clinical trials.10,11 Examples include a PEGylated peptidic substrate LUM01510,12 and an irreversible covalent probe VGT-309,11,13 both targeting cathepsins. While fluorescent probes have already begun to demonstrate significant value for surgical guidance, there remains a need for further strategies to improve key parameters such as tumor-to-background signal ratio (TBR), tissue selectivity and overall signal half-life.

One strategy to increase circulation time and tumor retention resulting in enhanced TBR involves increasing molecular weight and hydrophilicity. The use of PEGylation to in-crease probe size is a common design principle that lever-ages the enhanced permeability and retention (EPR) effect commonly observed in malignant tissues.14,15 One of the first enzyme-activated optical probes, introduced by Weissleder et al.,16 was a self-quenched PEGylated poly-lysine copolymer with molecular weight close to 500 kDa. The cathepsin-activated probe LUM015, currently under clinical investigation, also exploits a large PEG-based scaffold. We hypothesized that a hydrophilic, inert, and biocompatible macromolecular backbone, combined with an activity-based fluorescent substrate, could yield a probe with enhanced performance compared to its small-molecular counterparts. Furthermore, the macromolecular scaffold can be used to attach additional affinity-based targeting elements and to control the overall location and stoichiometry of quencher-fluorophore pairs. A recently developed class of macromolecular antibody mimetics, called iBodies, employs a plat-form based on N-(2-hydroxypropyl)methacrylamide copolymer (pHPMA).17 The concept of iBodies is based on facile derivatization in the polymer side chain with high-affinity ligands or inhibitors, fluorescent dyes and/or appropriate affinity tags, such as biotin, all on the same pHPMA carrier. Modular iBodies are similar to an antibody with high affinity, but with increased overall functionality and design flexibility similar to synthetic small molecules. iBody conjugates can be used for both imaging and inhibition of enzymes, and the pHPMA carrier enables control of density of displayed ligands to increase binding affinity and inhibi-tory potency.18–22 This platform has also been applied for specific binding of His-tagged proteins.23 Based on the success of iBodies, we hypothesized that pHPMA could be a promising template for improving the biodistribution and contrast of low-molecular weight protease-activated contrast agents. In this study, we describe macromolecular quenched fluorescent probes based on the pHPMA. We used this carrier to investigate the effects of quencher-fluorophore positioning and stoichiometry, as well as link-er choice, on overall probe performance in a mouse model of breast cancer. The pHPMA probes showed superior performance and biodistribution compared to the reference small molecular contrast agent. Furthermore, we found that their performance was dependent on the choice of linker and the structure of individual probes, providing a basis for the design and optimization of polymer-based quenched probes for targeting cancer and other diseases.

## RESULTS

Design of the macromolecular probes and their pre-cursors. We proposed that macromolecular probes derived from “iBodies” (Fig. 1a) would offer distinct advantages over their small-molecule counterparts by improving over-all biodistribution while leveraging the synthetic flexibility of a biocompatible hydrophilic backbone. The pHPMA precursors P1-P3 (Table S1) with controlled molecular weight and low dispersity were prepared by reversible addition-fragmentation chain transfer (RAFT) copolymerization of HPMA and 3-(3-methacrylamidopropanoyl)thiazolidine-2-thione (Ma-β-Ala-TT). The thiazolidine-2-thione (TT) reactive groups along the pHPMA chain enable attachment of amine terminated low-molecular weight contrast agents. Because multiple different components can be built into the pHPMA at controlled ratios, it is possible to regulate abundancy of each moiety on the carrier, allowing fine-tuning to optimize overall signal intensity. As a starting point for this study, we chose a previously published quenched pan-cathepsin substrate 6QC, containing sulfo-Cy5 as the fluorophore and sulfo-QSY21 as the quencher (Fig. 1b).24 Cathepsin-targeted probes have proven to be valuable tools for tumor imaging12,13,25–27 and 6QC, in particular, has been optimized for this application.24,28,29 We designed analogs of 6QC that introduce a spacer and a terminal amino group that can be used for incorporation onto the pHPMA precursor (Fig. 1d, 1f, 1h). We have previous-ly shown that the N-terminal benzyloxycarbonyl (Cbz) group plays an important role in the substrate recognition of 6QC, 27 therefore, we substituted the Cbz group with phenylglycine to preserve this motif in the spacer-modified analogs, which enabled the N-terminus to serve as a handle for spacer attachment. We designed several “architectures” available on the macromolecular carrier. In the primary probe design, we preserved the quencher-fluorophore arrangement of the original 6QC fragment where the two are positioned on the substrate peptide backbone (Fig. 1d). In case of the QF probe, protease cleavage releases the fluorophore from the pHPMA carrier as a free amine which can be protonated and retained in the acidic lysosomes of macro-phages (Fig. 1c). We then inverted the positions of the fluorophore and the quencher to generate the FQ probe (Fig. 1e, 1f). This configuration results in release of the quencher by the protease, leaving the fluorophore tethered to the pHPMA carrier. In addition, the tunable iBodies-based platform enables separation of the quencher and the fluorophore by tethering the two components to the pHPMA carrier separately (Fig. 1g). In this case, we attached the quencher to the pHPMA via a short non-cleavable link-er, while the fluorophore was attached using the cathepsin-cleavable substrate fragment (Fig. 1h). The iBody platform also enables precise control over the quencher-fluorophore ratio during synthesis. In this way, we can investigate how a change in the ratio of quenchers and fluorophores can affect the imaging properties of the probes.

**Figure 1.**
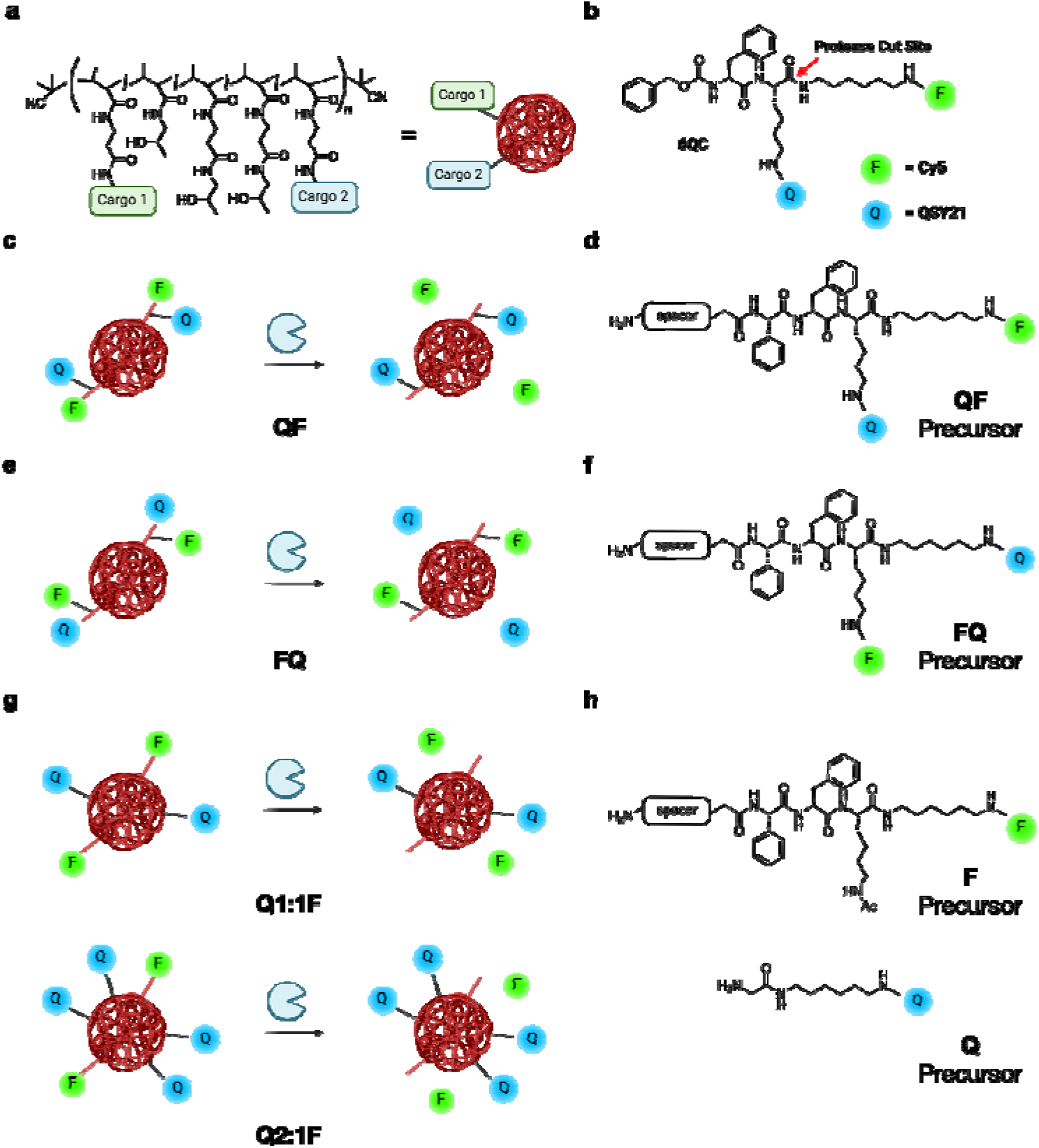
Chemical structure of employed *N*-(2-Hydroxypropyl)methacrylamide copolymer (pHPMA) and probe arrangements (a) Schematic representation of iBodies. Copolymer side chains are modified with various functional moieties via amide coupling (Cargo 1 and Cargo 2). Abundance and diversit of individual moieties can be controlled during pHPMA precursor modification. (b) Structure of the parental small molecule probe 6QC with protease cut site highlighted by red arrow. In this study, Q (blue sphere) represents sulfo-QSY21 while F (green sphere) represents sulfo-Cy5. (c-g) Proposed arrangements of probe on the polymer or “architectures” (c) The QF architecture where cleavage releases the fluorophore from the polymer, (d) Structure of the ligand used in the QF architecture, (e) The FQ architecture where cleavage releases the quencher from the polymer, (f) Structure of the FQ ligand, (g) Q1:1F and Q2:1F structures where quencher and fluorophore are attached separately at controlled ratios and cleavage releases the fluorophore from the polymer, (h) Structure of Q and F ligands for the Qx:1F architecture. Created with BioRender.com.

For our initial studies, we generated pHPMA probes with one-to-one (Q1:1F), two-to-one (Q2:1F) and three-to-one (Q3:1F) quencher-fluorophore ratios. Building upon this design, we synthesized a set of pHPMA probes to investigate the impact of the spacer and the architecture on probe performance.

In vitro evaluation of probe cleavage by recombinant cathepsin L and in RAW264.7 mouse macrophages. We employed three different linkers: short and flexible polyethylene glycol linker (PEG-4, referred to as short, S), a more rigid polyproline linker (13 prolines, polyPro, P) and a longer version of the polyethylene glycol linker (PEG-12, referred to as long, L) (Fig. 2a). The detailed compositions and characteristics of the pHPMA probes can be found in Table S2. We assessed the efficiency of purified cathepsin L (CatL) to cleave our macromolecular probes at two different probe concentrations (10 μM, 2.5 μM; to reduce influence of concentration-related effects such as aggregation of the macromolecular conjugates; Table S3). These results confirmed that the cleavage rate was highly dependent on linker length and only marginally influenced by probe architecture (Fig. 2b). Probes S-QF, S-Q1:1F, S-Q3:1F were cleaved at a slower rate than 6QC at both 10 μM and 2.5 μM. The polyPro probe P-QF was cleaved more efficiently than the S-QF probe with the initial cleavage rate at 2.5 μM almost reaching that of 6QC. Probes L-QF, L-FQ, L-Q1:1F, L-Q2:1F had the highest cleavage rates, exceeding that of 6QC at 2.5 μM, and reaching a relative rate of 2.20 ± 0.07 for L-Q2:1F (Table S3), suggesting that the rigidity of the polyPro linker did not yield significant benefit compared to the flexible PEG-12. All pHPMA probes were cleaved at a higher relative rate at the lower probe concentration 2.5 μM. We also analyzed the maximal signal-to-background ratio (SBR) of individual probes in the assay (Fig. 2c). This value is dependent on both the cleavage efficiency as well as the quenching efficiency, which is influenced by the quencher-fluorophore distance and ratio on the probe. For all the polymeric probes, we used fluorophore loadings of approximately two units per polymer chain (Table S2). The short PEG series probe S-QF had lower SBR than P-QF or L-QF, due to less efficient cleavage.

**Figure 2.**
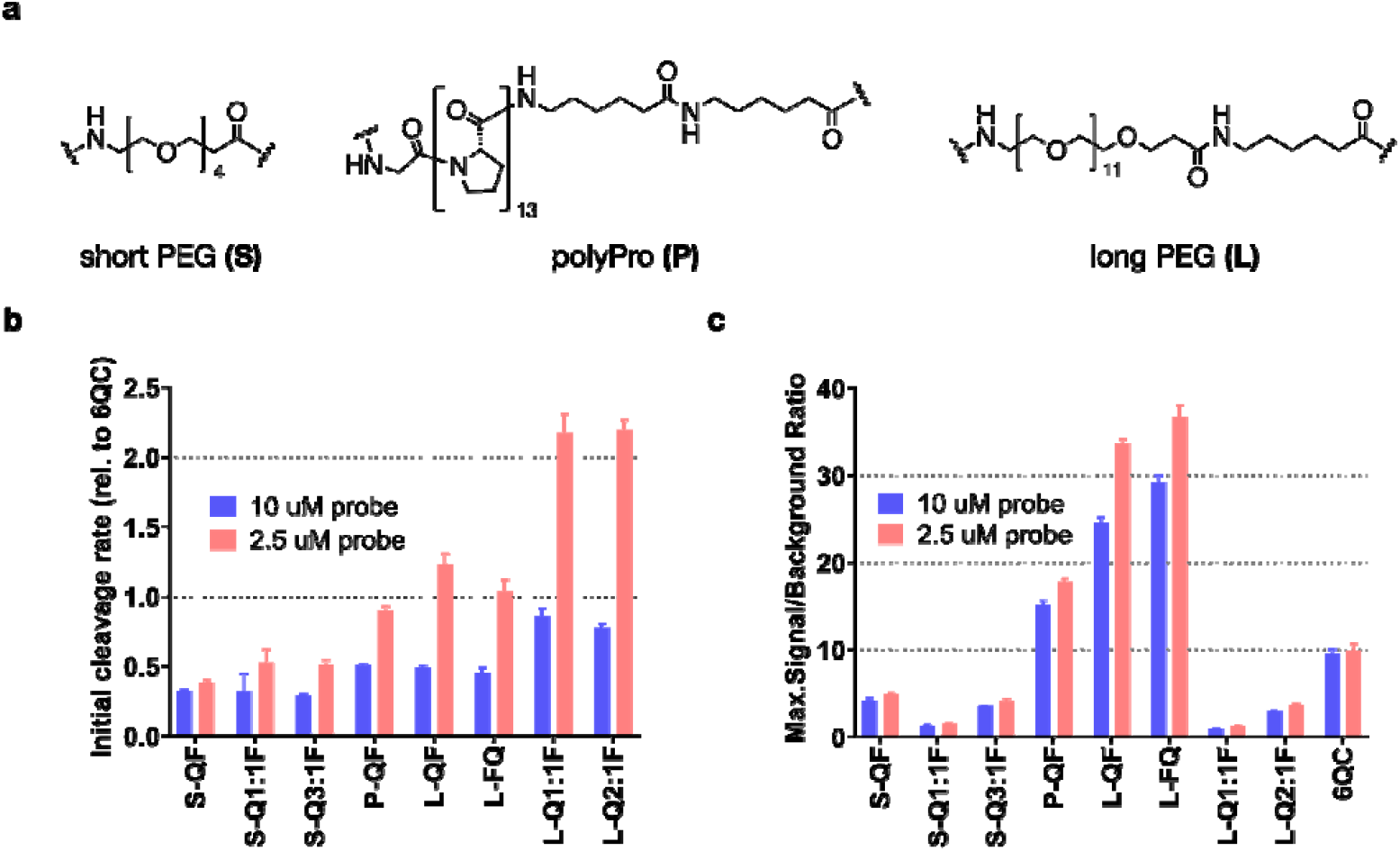
Effect of linker and architecture on *in vitro* probe activation by cathepsin L (CatL) (a) Structures of employed linkers. Polymer compositions and characteristics can be found in **Table S2**. (b) Bar graph showing initial cleavage rate of selected probes by purified cathepsin L. Rates were measured at 2 probe concentrations [10 μM, 2.5 μM] and normalized to 6QC as a standard. (c) Bar graph showing maximal signal-to-background ratio (SBR) calculated from the cleavage assay as a ratio between endpoint signal (90 min) in positive (CatL) and negative (no CatL) samples. SBR was measured at 2 concentrations [10 μM, 2.5 μM]. All experiments were performed in 96-well plates in triplicates. Data were collected over 90 min using Biotek Cytation 3 plate reader. Error bars represent standard deviation. All values can be found in **Table S3**.

However, we also observed a further drop in SBR values for S-Q1:1F and S-Q3:1F probes that bear quencher on the backbone (all values can be found in Table S3). This is likely due to increased quencher-fluorophore distance when the quencher is positioned on the pHPMA backbone. Swapping the position of sulfo-Cy5 and sulfo-QSY21 in the L-FQ probe did not influence the cleavage rate, nor the quenching efficiency significantly. Analogous to the short linker probes, the long linker probes L-Q1:1F, L-Q2:1F exhibited lower SBR than L-QF. Of all the probes tested, L-QF and L-FQ reached the highest SBR values, with an over three-fold increase compared to 6QC, likely due to efficient cleavage, quenching and the presence of multiple fluorophores on the pHPMA carrier. The Q1:1F architecture showed the lowest SBR in the individual series, while the addition of quencher moieties to the polymer led to an increase in SBR but never reached the values observed with QF/FQ architecture.

To monitor internalization and cleavage of selected pHPMA probes in comparison with the reference 6QC system in a cellular assay, we incubated the probes with RAW 264.7 mouse macrophages for 2 hours and recorded images using confocal microscopy before and after washing the cells with buffer (Fig. 3a). In the case of 6QC, strong specific signal was observed inside the cells after probe processing with no significant difference between the images before and after the wash. The pHPMA probes L-QF and L-FQ exhibited similar behavior to 6QC with limited signal present in solution and specific signal inside the cells. Generally, both polymer probes showed lower fluorescence intensity than 6QC after 2-hour incubation and there was no noticeable difference based on the quencher-fluorophore arrangement. Furthermore, we included a non-quenched control probe (nQ) in which the fluorophore was directly tethered to the pHPMA via a short non-cleavable linker. For the nQ probe, strong fluorescent signal was observed in solution after incubation, as highlighted by the difference between the images before and after the wash. This was further con-firmed by analysis of Cy5 fluorescence intensity in the medium (Fig. 3b). Nevertheless, probe nQ was internalized by RAW 264.7 macrophages and could be detected inside the cells after washing. Probe L-Q2:1F generated similar intensity of fluorescence inside the cells after the wash as the nQ probe with overall reduced background fluorescence in the media (Fig. 3a, 3b).

**Figure 3.**
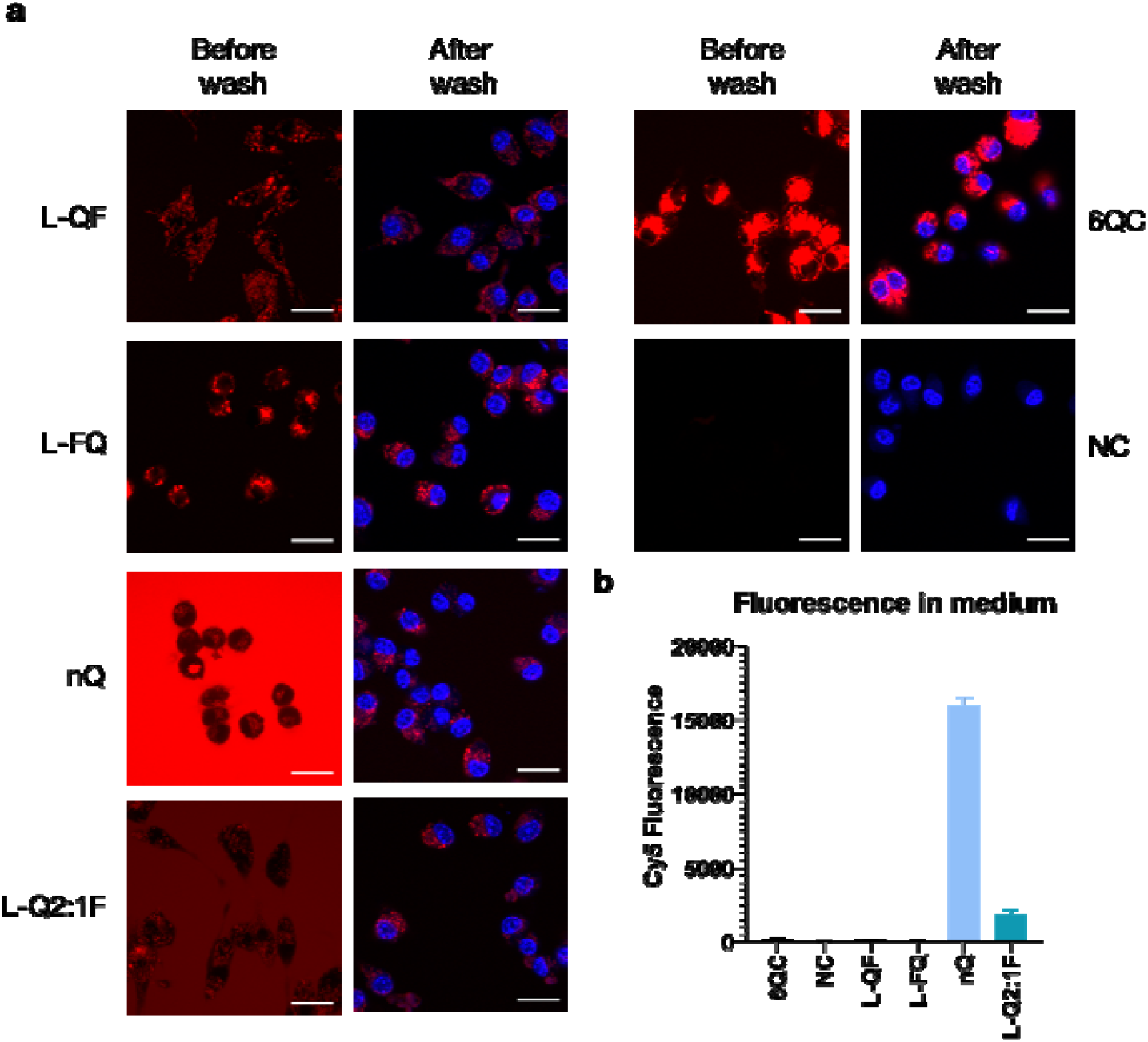
Internalization and cleavage of probes in RAW 264.7 mouse macrophages (a) Confocal microscopy images of RAW 264.7 cells incubated with the indicated probes for 2 hours at 37 °C. Cells were imaged directly after 2 hours (first column) or washed and stained with Hoechst 33342 (second column). Imaging was performed with excitation at 639 nm and emission at 669 nm for Cy5 (colored red) and excitation at 405 nm and emission at 435 nm for Hoechst (colored blue). Scale bar = 20 μm. (b) Bar graph showing average Cy5 fluorescence intensity in conditioned medium after 2 hours of incubation. Data is present for four experiments. Error bars represent standard deviation.

Evaluation of probe performance in vivo in a mouse model. We assessed the performance of the probes in vivo using an orthotopic 4T1 triple negative breast cancer mouse model. For these experiments, mice were subcutaneously injected with 4T1 cancer cells. After one week of tumor growth, the probes were intravenously administered retro-orbitally. Images were acquired at multiple timepoints to examine the evolution of fluorescent signal intensity and the tumor-to-background ratio (TBR) over time (Fig. 4a). All quenched probes accumulated at the tumor site, with similar efficiency as the reference probe, 6QC. The P-QF probe showed increased background fluorescent signal in the abdominal area, and the non-quenched probe nQ exhibited high fluorescent signal in the surrounding tissue. Next, we determined the TBRs of all probes at three different timepoints (2h, 8h, 24h; Fig. 4b). The first three macro-molecular probes (S-QF, P-QF, L-QF) all possess a QF architecture, differing only in the linker structure. General-ly, TBRs of all QF architecture probes were stable over time. The P-QF and L-QF probes exhibited significantly higher TBRs than 6QC at all studied timepoints, while S-QF showed significantly higher TBR than 6QC only at the 2h timepoint. At the 24h timepoint, P-QF and L-QF reached TBRs of 2.9 ± 0.4 and 3.1 ± 0.8, respectively, compared to 2.3 ± 0.5 for 6QC. The differences between P-QF and L-QF were generally small and statistically insignificant. However, the poly proline probe P-QF showed increased accumulation in kidneys and liver compared to the PEG-based probes (Fig. S1), therefore, we prioritized the PEG-12 linker (L), which performed well in both in vitro and in vivo evaluation, to assess architecture influence on probe performance.

**Figure 4:**
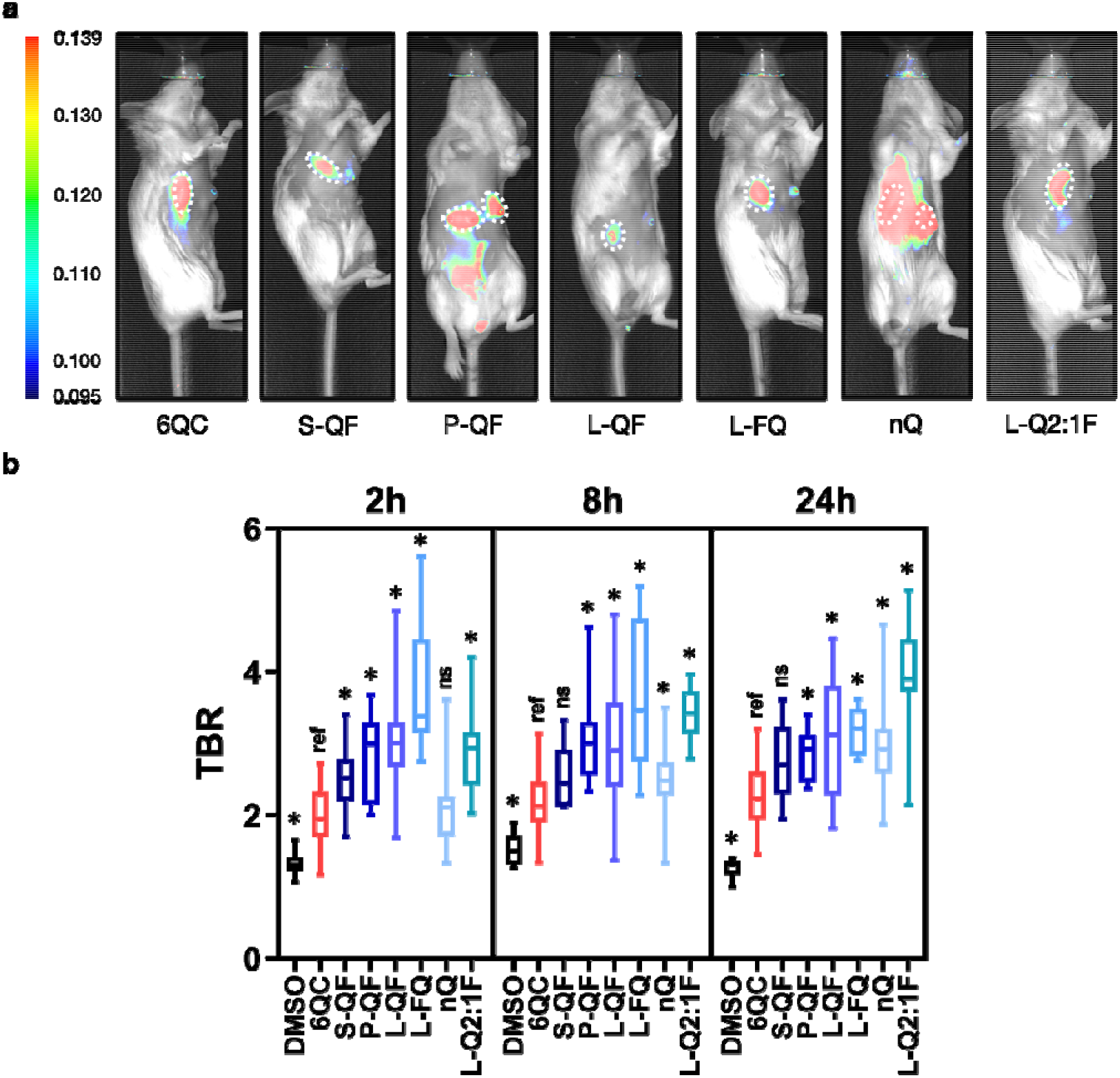
*In vivo* Probe Evaluation: Quantification of Tumor-to-Background Ratio (TBR) for individual probes in live mice (a) Representative fluorescence images showing a mouse with two breast tumors after injection of indicated probe [6.25 nmol] 24 h prior to imaging. Tumors are highlighted using a white dotted line. The color bar shows fluorescence intensity in arbitrary units. (b) Box plots showing tumor-to-background ratio (TBR) quantification of selected probes in live mice after 2, 8 and 24 hours. *N:* 3 to 10 mice, 6 to 20 tumors per condition. Images were acquired using the Pearl Trilogy small animal imaging system at the indicated timepoints. Statistics were calculated via the Wilcoxon rank-sum test, asterisk represents statistical significance (p< 0.05), ns – not significant. Boxes extend from the 25^th^ to the 75^th^ percentile with whiskers extending to minimal and maximal values. The line in the middle of the box is plotted at the median value for each sample set. All values can be found in **Table S4**.

We introduced the flipped architecture L-FQ, which re-leases the quencher instead of the fluorophore, and L-Q2:1F as the representative of the Qx:1F architecture. Similar to L-QF, the L-FQ probe exhibited significantly high-er TBR than 6QC at all analyzed timepoints (Fig. 4b). The change in the quencher-fluorophore arrangement did not influence TBR significantly. Interestingly, the nQ probe (TBR = 3.0 ± 0.8, 24h), driven solely by the EPR effect, also provided high contrast, but required 8 hours to establish the significant difference from 6QC and 24 hours to achieve peak TBR values. The L-Q2:1F probe also showed increasing TBR values over time and reached the highest TBR (3.9 ± 0.9, 24h) of all the probes tested.

To accurately assess the overall tumor tissue specificity of the probes, we also imaged sacrificed mice after the final timepoint with their skin splayed open to reveal the under-surface of the tumors (Fig. 5a). Similar to what was observed in live mice, all quenched probes labeled tumors effectively, mostly varying in signal intensity and back-ground fluorescence in the non-tumor tissues. The QF architecture probes exhibited TBRs ranging from 3.9 ± 1.5 (L-QF) to 5.1 ± 1.9 (S-QF), compared to 2.3 ± 0.4 of 6QC (Fig. 5b). Interestingly, S-QF showed the highest contrast of the QF probes in this configuration. Overall, TBR values were in good correlation with the data from live mice. The L-Q2:1F (TBR = 5.1 ± 1.3) and S-QF (TBR = 5.1 ± 1.9) probes achieved highest TBR values in this configuration, while L-FQ (TBR = 3.4 ± 1.0) and nQ (TBR = 3.7 ± 1.4) showed slightly lower contrast com-pared to other polymeric probes. Furthermore, we analyzed biodistribution of the probes by imaging harvested organs. Accumulation in kidneys and lungs was significantly reduced for all macromolecular probes compared to 6QC, while most of them also exhibited significantly reduced accumulation in heart (Figure S1). Interestingly, for most quenched probes, liver accumulation was not significantly different from that of 6QC, while S-QF and nQ showed significant reduction in liver signals.

**Figure 5.**
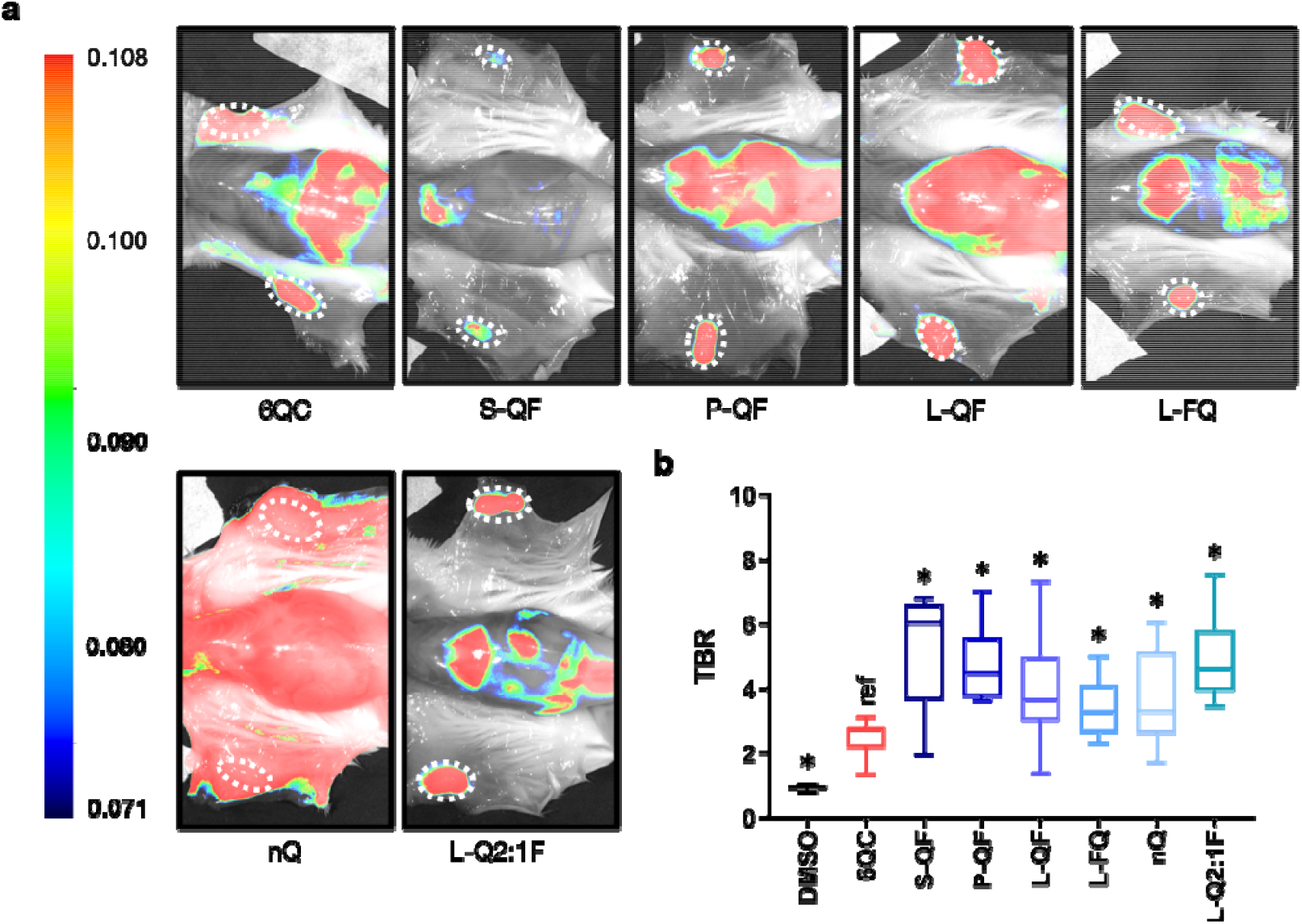
*In vivo* Probe Evaluation: Quantification of Tumor-to-Background Ratio (TBR) for individual probes in splayed mice (a) Representative fluorescence images showing a splayed mouse with two tumors after injection of indicated probe [6.25 nmol] 24 h prior to imaging. Tumors are highlighted using white dotted lines. Color bar shows fluorescence intensity in arbitrary units. (b) Box plot showing tumor-to-background ratio (TBR) quantification of selected probes in splayed mice. *N:* 3 to 10 mice, 6 to 20 tumors per condition. Images were acquired using the Pearl Trilogy small animal imaging system. Statistics were calculated via the Wilcoxon rank-sum test, asterisk represents statistical significance (p< 0.05). Boxes extend from the 25th to the 75th percentile with whiskers extending to minimal and maximal values. The line in the middle of the box is plotted at the median value for each data set. All values can be found in **Table S4**.

## DISCUSSION

Optimizing signal-to-background ratio and increasing tis-sue specificity remain important goals in probe development. This can be achieved via choice of appropriate activation mechanism, optimization of activation efficiency and finally regulation of overall pharmacodynamics of a probe. We aimed to optimize a small molecule substrate probe by transferring it to a macromolecular core scaffold which would allow control over probe size, positioning of fluorophore and quencher, and density of ligands on the back-bone. In this study, we investigated several types of flexible or rigid linkers of varying lengths to identify an optimal configuration for cathepsin cleavage. We then investigated positioning as well as ratios of the quenchers and the fluorophores. This allowed us to confirm that the HPMA copolymer backbone is a valuable tool for enhancing properties of low molecular weight quenched fluorescent probes.

Specifically, we found that linker length is important for efficient probe cleavage, while it may also affect probe bio-distribution. Our data suggest that relatively long linkers are required to allow efficient cleavage by the appropriate protease, probably due to the large size of the polymer backbone (Fig. 2b). The rigidness of the polyPro linker did not provide an advantage over the flexible PEG-12 (Fig. 2b). Interestingly, the P-QF probe also showed increased kidney accumulation compared to the PEG-based S-QF and L-QF (Fig. S1). The length of the linker also has an im-pact on the distance between the fluorophore and the quencher in Qx:1F architectures, where the quencher is located on the polymer backbone (Fig. 2c). Therefore, it is important to carefully select the linker length to achieve a balance between cleavage and quenching efficiency.

The data from cell culture models matched the results observed in the purified enzyme assay (Fig. 3). Probes L-QF and L-FQ showed high signal-to-background ratio due to efficient quenching. The lower signal intensity at the 2-hour timepoint for the polymeric probes compared to the reference probe 6QC is likely caused by slower internalization of the large polymer molecules. Interestingly, there was no significant difference in signal retention between probe L-QF, which releases the fluorophore, and probe L-FQ, which releases the quencher. This suggests that both the dye linked to the polymer and the free dye are retained inside lysosomes, resulting in a durable fluorescent signal over time for both probe architectures. Probes L-Q2:1F and nQ also produced signal inside the cells but exhibited higher fluorescence in the medium. This high background due to lack of quenching is likely translated into lower TBR values for these probes in vivo, especially at early timepoints, before free probes are cleared from the bloodstream.

Our in vivo experiments showed that, regardless of linker and probe structure, use of a polymer backbone improved probe biodistribution and contrast compared to the small molecule probe 6QC. We also found the trends of TBR values in vivo correlated with the trends in enzyme cleavage rates in vitro, even though the TBR differences were subtle. An unexpected observation was that the S-QF probe, which did not show superior contrast compared to other probes in live mice, exhibited the highest TBR in the splay setup (Fig. 5b). This could potentially be explained by increased overall non-specific tissue accumulation of the longer, more lipophilic linkers. Another interesting trend observed was that the iBody probes generally reached higher TBR values in the splay setup, while the values for 6QC did not show a significant difference between the live mice and the splayed mice (Fig. 4b, 5b). The most striking observation arises from the probe L-Q2:1F that showed poor quenching efficiency in vitro (Fig. 2c) but, surprisingly, achieved the highest TBR in vivo in live mice in our dataset (Fig. 4b). This may be in part explained by the significant contribution of the EPR effect.14,15 The nQ probe, which is driven solely by this effect, reached TBR values superior to 6QC, but only at the late timepoints due to high background after injection (Fig. 4b). Our results demonstrate that quenching, even if only partial, can act synergistically with the enhanced uptake effects to dramatically improve contrast. Even though probe L-Q2:1F exhibited higher TBR values in live mice compared to the other probes, it still needs up to 24 hours to develop this contrast (Fig. 4b). The slow generation of contrast appears to be specific to non- or poorly quenched probes, as it is a common trait for nQ and L-Q2:1F. In contrast, probes with high quenching efficiency, such as L-QF, reach their peak TBR values as early as 2 hours after application. This rapid signal accumulation may be more compatible with common surgical workflows.

Overall, the iBody approach provides high modularity and fine-tuning of probe signal intensity, contrast, and kinetics. In the future, we plan to include affinity-based targeting elements to the polymer backbone to further im-prove biodistribution and to direct conjugates to specific tissues or cell types.

Possible targets could include fibro-blast activation protein (FAP),30 glutamate carboxypeptidase II (GCPII)31 or folate receptor α (FRα).9 This scaffold is also ideally suited for AND-gate type probes reported in the literature that rely on subsequent cleavage of two orthogonal protease sequences, which increases specificity and sensitivity dramatically.32 AND-gate probes eliminate false positive signal caused by serendipitous cleavage of the single substrate sequence in a non-target tissue and could help to reduce the relatively high fluorescent signal in the liver and kidneys. Finally, the flexible iBody approach is also well suited for ratiometric imaging approaches that can eliminate false positive signals caused by local accumulation of contrast agents.33 While the EPR effect helps to increase tumor tissue accumulation of macromolecular probes, similar effects can cause probe uptake in unwanted areas, resulting in a false positive signal and, therefore, increase the background of the probe. The ratiometric approach applied to the probes described here should help minimize this effect, resulting in enhanced contrast. Other modifications, such as the incorporation of clinically relevant chromophores, for example, ICG or IRDye800CW,3 can also be applied and should not influence the physicochemical properties and biodistribution dramatically due to the primary contribution of the polymer backbone.

## CONCLUSIONS

This work demonstrates that polymers based on the N-(2-hydroxypropyl)methacrylamide copolymer (pHPMA) pro-vide and optimal scaffold for use in the design of optical imaging probes. The simple and modular synthesis of these polymers enable engineering of overall ligand density as well as architecture of dye and quenchers on the probe. Furthermore, this strategy enables a flexible strategy for functionalization of the probes using multiple targeting ligands. We show here that pHMPA-based probes containing a cathepsin cleavable substrate have improved properties for imaging of tumor margins during surgery compared to simple peptide scaffolds. These improvements include in-creased fluorescent signal and enhanced overall contrast. We therefore believe this overall approach using the pHMPA scaffold is likely to be valuable for the design and synthesis of a wide range of contrast agents for imaging cancer or other relevant markers of human diseases.

## EXPERIMENTAL SECTION

Synthesis of Target Linker-Modified Probe Derivatives, Monomers, Copolymer Precursors and Copolymer Conjugates. Target linker-modified probe derivatives were prepared by a combination of solid-phase and liquid-phase peptide chemistry. Monomers were prepared using standard procedures available in the literature. Copolymer precursors were prepared by RAFT copolymerization and conjugated with ligands of interest via aminolytic reaction of polymer precursor containing TT reactive groups. De-tailed description of materials, procedures and characterizations can be found in the Supplementary Information (SI).

Assay for enzymatic cleavage of probes by recombinant cathepsin L. Buffer for enzymatic assay was prepared by dissolving citrate (50 mM), Triton X-100 (0.1 %), CHAPS (0.5 %) in MiliQ water and the pH was adjusted to 5.5. DTT (0.8 mg/mL) was added freshly before use. The assay was performed in a 96-well plate. Positive samples were prepared by mixing enzyme (2.2 nM, 45 μL) in assay buffer and substrate (100 μM or 25 μM, 5 μL) dis-solved in MiliQ water. Negative samples were prepared by mixing assay buffer (45 μL) and substrate (100 μM or 25 μM, 5 μL). The assay was performed at 37 °C. Fluorescence values were detected by a plate reader (exc. 640 nm, em. 670 nm) every 45 s over 90 min. All experiments were performed in triplicates. Initial cleavage rate was evaluated as slope of the regression curve to the initial linear phase of the kinetic curve (first 12 points). Signal-to-background ratio (SBR) was evaluated as ratio between fluorescence intensity of the positive sample at 90 min and mean fluorescence intensity of negative samples at 90 min.

General Cell Culture Methods. All cell lines were passaged a minimum of three times after thawing before use in confocal microscopy experiments and before injections into mice. Both 4T1 cells (ATCC CRL-2539) and RAW246.7 macrophages (ATCC TIB-71) were grown with 100 U/mL penicillin and 100□μg/mL streptomycin and with 10% Fetal Animal Serum FAS supplemented into the media.

4T1 cells were cultured in Roswell Park Memorial Institute (RPMI, Corning, 10-040-CV) 1640 medium containing 2□g/L of glucose, 0.3□g/mL of L-glutamine. RAW246.7 mouse macrophages were cultured in Dulbecco’s Modified Eagle’s Medium (ATCC, 30-2002) with 10% FAS.

Confocal Microscopy. RAW246.7 cells were distributed on 4-chamber microscopy dishes (Cellvis, D35C4201.5N). Next day, cells were washed, and probes were added in phenol-red free DMEM (Gibco, 21063029), 10% FAS. After incubation at 37 °C, 5% CO2, 2 hours, cells were either directly imaged or washed 1x and incubated for 5 min with Hoechst 33342 (Tocris, 5117/50) to a final concentration 1 ng/µL prior to imaging. Images were acquired using a Zeiss LSM700 fluorescence confocal microscope with 405 nm laser for Hoechst 33342 staining and 639 nm for probe staining with 63x magnification (Plan-Apochromat 63x/1.40 Oil DIC M27). All settings (laser %, gain, pinhole) were set at the beginning of the imaging and then kept constant throughout. Experiment was performed more than three times and representative images were selected for the final figure.

4T1 Breast Tumor Model. 4T1 cells were prepared according to the procedure described in the SI. While under isoflurane anesthesia, mice were subcutaneously injected in the third and eighth mammary fat pads of BALB/c female mice (aged 1–8 weeks; Jackson Laboratory) with 100 μL of the diluted 4T1 cells (1□×□105 cells per fat pad). Once the tumors were developed, mice were injected intravenously with 6.25 nmol of probe in 100 μL of injection solution using a 28-gauge 1 mL insulin syringe into the tail vein. Post injection, the mice were non-invasively imaged using the LI-COR Pearl Trilogy imaging system at multiple time points. After the live mice imaging, mice were euthanized using cervical dislocation under isoflurane anesthesia. Mice were then splayed and imaged. Finally, through dissection, organs were collected (liver, kidneys, spleen, lungs, heart), as well as the primary tumors and non-injected fat pad controls. These tissues were then imaged ex-vivo. Fluorescence intensity was measured using the built-in LI-COR Image Studio software. Additional information regarding probe formulation, imaging conditions and evaluation of obtained images can be found in the SI. All images shown were linked to display the same brightness and contrast settings for a given condition (example: 24-hour images for all probes are linked).

Statistical Analysis. Statistical analysis was performed in Excel (Microsoft, Redmond, WA, USA) and Prism (GraphPad Software, San Diego, CA, USA). Statistical significance was calculated based on the Wilcoxon rank-sum test and the Brown-Forsythe and Welch ANOVA test.

## ACKNOWLEDGMENT

Research reported in this publication was supported by the National Institutes of Health under award number R01EB028628 (to M.B.), the project National Institute for Cancer Research (Programme EXCELES, ID Project No. LX22NPO5102) funded by the European Union - Next Generation EU, the European Social Fund - Operational Pro-gramme Research, Development and Education (Project “IOCB Mobility II”, No. CZ.02.2.69/0.0/0.0/18_053/0016940), the Ministry of Health of Czech Republic project (NU21-08-00280), and the Czech Science Foundation (Project No. 23-05642S). The authors thank Jitka Bařinková for her technical assistance, Dominik Musil for his help with the in vitro assay, and Kateřina Radilová for help with visually improving the figures.

## ABBREVIATIONS

FDA: the U.S. Food and Drug Administration
NIR: near-infrared
FRα: folate receptor α
TBR: tumor-to-background ratio
EPR: enhanced permeability and retention
pHPMA: N-(2-hydroxypropyl)methacrylamide copolymer
RAFT: reversible addition-fragmentation chain transfer
TT: thiazolidine-2-thione
Cbz: benzyloxycarbonyl
PEG: polyethylene glycol
CatL: cathepsin L
SBR: signal-to-background ratio
FAP: fibroblast activation protein
GCPII: glutamate carboxypeptidase II
min: minutes

## Notes

### Competing Interest Statement

The authors have declared no competing interest.

## REFERENCES

1. Barth, C. W.; Gibbs, S. Fluorescence Image-Guided Surgery: A Perspective on Contrast Agent Development. In Molecular-Guided Surgery: Molecules, Devices, and Applica-ions VI; Proc. SPIE 11222, 2020, p 18.

2. Garland, M.; Yim, J. J.; Bogyo, M. A Bright Future for Precision Medicine: Advances in Fluorescent Chemical Probe Design and Their Clinical Application. Cell. Chem. Biol. 2016, 23 (1), 122–136.

3. Seah, D.; Cheng, Z.; Vendrell, M. Fluorescent Probes for Imaging in Humans: Where Are We Now? ACS Nano 2023, 17, 19478–19490.

4. Olson, M. T.; Ly, Q. P.; Mohs, A. M. Fluorescence Guidance in Surgical Oncology: Challenges, Opportunities, and Translation. Mol. Imaging Biol. 2018, 21 (2), 200–218.

5. Hong, G.; Antaris, A. L.; Dai, H. Near-Infrared Fluor-ophores for Biomedical Imaging. Nat. Biomed. Eng. 2017 1:1 2017, 1 (1), 1–22.

6. Peltrini, R.; Podda, M.; Castiglioni, S.; Di Nuzzo, M. M.; D’Ambra, M.; Lionetti, R.; Sodo, M.; Luglio, G.; Mucilli, F.; Di Saverio, S.; Bracale, U.; Corcione, F. Intraoperative Use of Indocyanine Green Fluorescence Imaging in Rectal Cancer Surgery: The State of the Art. World J. Gastroenterol. 2021, 27 (38), 6374.

7. Seitkazina, A.; Yang, J.-K.; Kim, S. Clinical Effectiveness and Prospects of Methylene Blue: A Systematic Re-view. Precis. Future Med. 2022, 6 (4), 193–208.

8. Hoogstins, C. E. S.; Tummers, Q. R. J. G.; Gaarenstroom, K. N.; de Kroon, C. D.; Trimbos, J. B. M. Z.; Bosse, T.; Smit, V. T. H. B. M.; Vuyk, J.; van de Velde, C. J. H.; Cohen, A. F.; Low, P. S.; Burggraaf, J.; Vahrmeijer, A. L. A Novel Tumor-Specific Agent for Intraoperative Near-Infrared Fluorescence Imaging: A Translational Study in Healthy Volunteers and Patients with Ovarian Cancer. Clin. Cancer Res. 2016, 22 (12), 2929–2938.

9. Tanyi, J. L.; Randall, L. M.; Chambers, S. K.; Butler, K. A.; Winer, I. S.; Langstraat, C. L.; Han, E. S.; Vahrmeijer, A. L.; Chon, H. S.; Morgan, M. A.; Powell, M. A.; Tseng, J. H.; Lopez, A. S.; Wenham, R. M. A Phase III Study of Pafolacianine Injection (OTL38) for Intraoperative Imaging of Folate Receptor-Positive Ovarian Cancer (Study 006). J. Clin. Oncol. 2023, 41 (2), 276–284.

10. Investigation of Novel Surgical Imaging for Tumor Excision (INSITE), NCT03686215, 2023. https://clinicaltrials.gov/study/NCT03686215 x(accessed 2024-03-29).

11. Phase 2 Study of VGT-309 in Lung Cancer, NCT05400226, 2023. https://classic.clinicaltrials.gov/ct2/show/NCT05400226 x(accessed 2024-03-29).

12. Whitley, M. J.; Cardona, D. M.; Lazarides, A. L.; Spasojevic, I.; Ferrer, J. M.; Cahill, J.; Lee, C. L.; Snuderl, M.; Blazer, D. G.; Hwang, E. S.; Greenup, R. A.; Mosca, P. J.; Mito, J. K.; Cuneo, K. C.; Larrier, N. A.; O’Reilly, E. K.; Riedel, R. F.; Eward, W. C.; Strasfeld, D. B.; Fukumura, D.; Jain, R. K.; Lee, W. D.; Griffith, L. G.; Bawendi, M. G.; Kirsch, D. G.; Brigman, B. E. A Mouse-Human Phase 1 Co-Clinical Trial of a Protease-Activated Fluorescent Probe for Imaging Cancer. Sci. Transl. Med. 2016, 8 (320), 320ra4.

13. Kennedy, G. T.; Holt, D. E.; Azari, F. S.; Bernstein, E.; Nadeem, B.; Chang, A.; Sullivan, N. T.; Segil, A.; Desphande, C.; Bensen, E.; Santini, J. T.; Kucharczuk, J. C.; Delikatny, E. J.; Bogyo, M.; Egan, A. J. M.; Bradley, C. W.; Eruslanov, E.; Lickliter, J. D.; Wright, G.; Singhal, S. A Cathepsin-Targeted Quenched Activity-Based Probe Facilitates Enhanced Detection of Human Tumors during Resection. Clin. Cancer Res. 2022, 28 (17), 3729–3741.

14. Maeda, H. Macromolecular Therapeutics in Cancer Treatment: The EPR Effect and Beyond. J. Control. Release 2012, 164 (2), 138–144.

15. Fang, J.; Nakamura, H.; Maeda, H. The EPR Effect: Unique Features of Tumor Blood Vessels for Drug Delivery, Factors Involved, and Limitations and Augmentation of the Effect. Adv. Drug Deliv. Rev. 2011, 63 (3), 136–151.

16. Weissleder, R.; Tung, C.-H.; Mahmood, U.; Bogdanov, A. In Vivo Imaging of Tumors with Protease-Activated near-Infrared Fluorescent Probes. Nat. Biotechnol. 1999, 17 (4), 375–378.

17. Šácha, P.; Knedlík, T.; Schimer, J.; Tykvart, J.; Pa-rolek, J.; Navrátil, V.; Dvořáková, P.; Sedlák, F.; Ulbrich, K.; Strohalm, J.; Majer, P.; Šubr, V.; Konvalinka, J. IBodies: Modular Synthetic Antibody Mimetics Based on Hydrophilic Pol-ymers Decorated with Functional Moieties. Angew. Chem. Int. Ed. 2016, 55 (7), 2356–2360.

18. Pospíšilová, K.; Knedlík, T.; Šácha, P.; Kostka, L.; Schimer, J.; Brynda, J.; Král, V.; Cígler, P.; Navrátil, V.; Etrych, T.; Šubr, V.; Kugler, M.; Fábry, M.; řezácová, P.; Konvalinka, J. Inhibitor– Polymer Conjugates as a Versatile Tool for Detection and Visualization of Cancer-Associated Carbonic Anhydrase Isoforms. ACS Omega 2019, 4 (4), 6746–6756.

19. Blažková, K.; Beranová, J.; Hradilek, M.; Kostka, L.; Šubr, V.; Etrych, T.; Šácha, P.; Konvalinka, J. The Development of a High-Affinity Conformation-Sensitive Antibody Mimetic Using a Biocompatible Copolymer Carrier (IBody). J. Biol. Chem. 2021, 297 (5), 101342.

20. Šimon, P.; Knedlík, T.; Blažková, K.; Dvořáková, P.; Březinová, A.; Kostka, L.; Šubr, V.; Konvalinka, J.; Šácha, P. Identification of Protein Targets of Bioactive Small Molecules Using Randomly Photomodified Probes. ACS Chem. Biol. 2018, 13 (12), 3333–3342.

21. Dvořáková, P.; Bušek, P.; Knedlík, T.; Schimer, J.; Etrych, T.; Kostka, L.; Stollinová Šromová, L.; Šubr, V.; Šácha, P.; Šedo, A.; Konvalinka, J. Inhibitor-Decorated Polymer Conjugates Targeting Fibroblast Activation Protein. J. Med. Chem. 2017, 60 (20), 8385–8393.

22. Šubr, V.; Ormsby, T.; Šácha, P.; Konvalinka, J.; Etrych, T.; Kostka, L. The Role of the Biotin Linker in Poly-mer Antibody Mimetics, IBodies, in Biochemical Assays. Polym. Chem. 2021, 12 (41), 6009–6021.

23. Beranová, J.; Knedlík, T.; Šimková, A.; Šubr, V.; Kostka, L.; Etrych, T.; Šácha, P.; Konvalinka, J. Tris-(Nitrilotriacetic Acid)-Decorated Polymer Conjugates as Tools for Immobilization and Visualization of His-Tagged Proteins. Catalysts 2019, 9 (12), 1011.

24. Ofori, L. O.; Withana, N. P.; Prestwood, T. R.; Verdoes, M.; Brady, J. J.; Winslow, M. M.; Sorger, J.; Bogyo, M. Design of Protease Activated Optical Contrast Agents That Exploit a Latent Lysosomotropic Effect for Use in Fluorescence-Guided Surgery. ACS Chem. Biol. 2015, 10 (9), 1977–1988.

25. Verdoes, M.; Oresic Bender, K.; Segal, E.; van der Linden, W. A.; Syed, S.; Withana, N. P.; Sanman, L. E.; Bogyo, M. Improved Quenched Fluorescent Probe for Imaging of Cysteine Cathepsin Activity. J. Am. Chem. Soc. 2013, 135 (39), 14726–14730.

26. Poreba, M.; Groborz, K.; Vizovisek, M.; Maruggi, M.; Turk, D.; Turk, B.; Powis, G.; Drag, M.; Salvesen, G. S. Fluorescent Probes towards Selective Cathepsin B Detection and Visualization in Cancer Cells and Patient Samples. Chem. Sci. 2019, 10 (36), 8461–8477.

27. Tholen, M.; Yim, J. J.; Groborz, K.; Yoo, E.; Martin, B. A.; van den Berg, N. S.; Drag, M.; Bogyo, M. Design of Optical Imaging Probes by Screening of Diverse Substrate Libraries Directly in Disease Tissue Extracts. Angew. Chem. Int. Ed. Engl. 2020, 59 (43), 19143–19152.

28. Yim, J. J.; Tholen, M.; Klaassen, A.; Sorger, J.; Bogyo, M. Optimization of a Protease Activated Probe for Optical Surgical Navigation. Mol. Pharm. 2018, 15 (3), 750–758.

29. Yim, J. J.; Harmsen, S.; Flisikowski, K.; Flisikow-ska, T.; Namkoong, H.; Garland, M.; van den Berg, N. S.; Vilches-Moure, J. G.; Schnieke, A.; Saur, D.; Glasl, S.; Gorpas, D.; Habtezion, A.; Ntziachristos, V.; Contag, C. H.; Gambhir, S. S.; Bogyo, M.; Rogalla, S. A Protease-Activated, near-Infrared Fluorescent Probe for Early Endoscopic Detection of Premalignant Gastrointestinal Lesions. Proc. Natl. Acad. Sci. U.S.A. 2021, 118 (1), e2008072118.

30. Roy, J.; Hettiarachchi, S. U.; Kaake, M.; Mukkamala, R.; Low, P. S. Design and Validation of Fibroblast Activation Protein Alpha Targeted Imaging and Therapeutic Agents. Theranostics 2020, 10 (13), 5778–5789.

31. A. Foss, C.; C. Mease, R.; Y. Cho, S.; J. Kim, H.; G. Pomper, M. GCPII Imaging and Cancer. Curr. Med. Chem. 2012, 19 (9), 1346–1359.

32. Widen, J. C.; Tholen, M.; Yim, J. J.; Antaris, A.; Casey, K. M.; Rogalla, S.; Klaassen, A.; Sorger, J.; Bogyo, M. AND-Gate Contrast Agents for Enhanced Fluorescence-Guided Surgery. Nat. Biomed. Eng. 2020, 5 (3), 264–277.

33. Faucher, F. F.; Liu, K. J.; Cosco, E. D.; Widen, J. C.; Sorger, J.; Guerra, M.; Bogyo, M. Protease Activated Probes for Real-Time Ratiometric Imaging of Solid Tumors. ACS Cent. Sci. 2023, 9 (5), 1059–1069.

