## Supporting Information for "Polymer-tethered quenched fluorescent probes for enhanced imaging of tumor associated proteases"

### Supplementary figures and tables

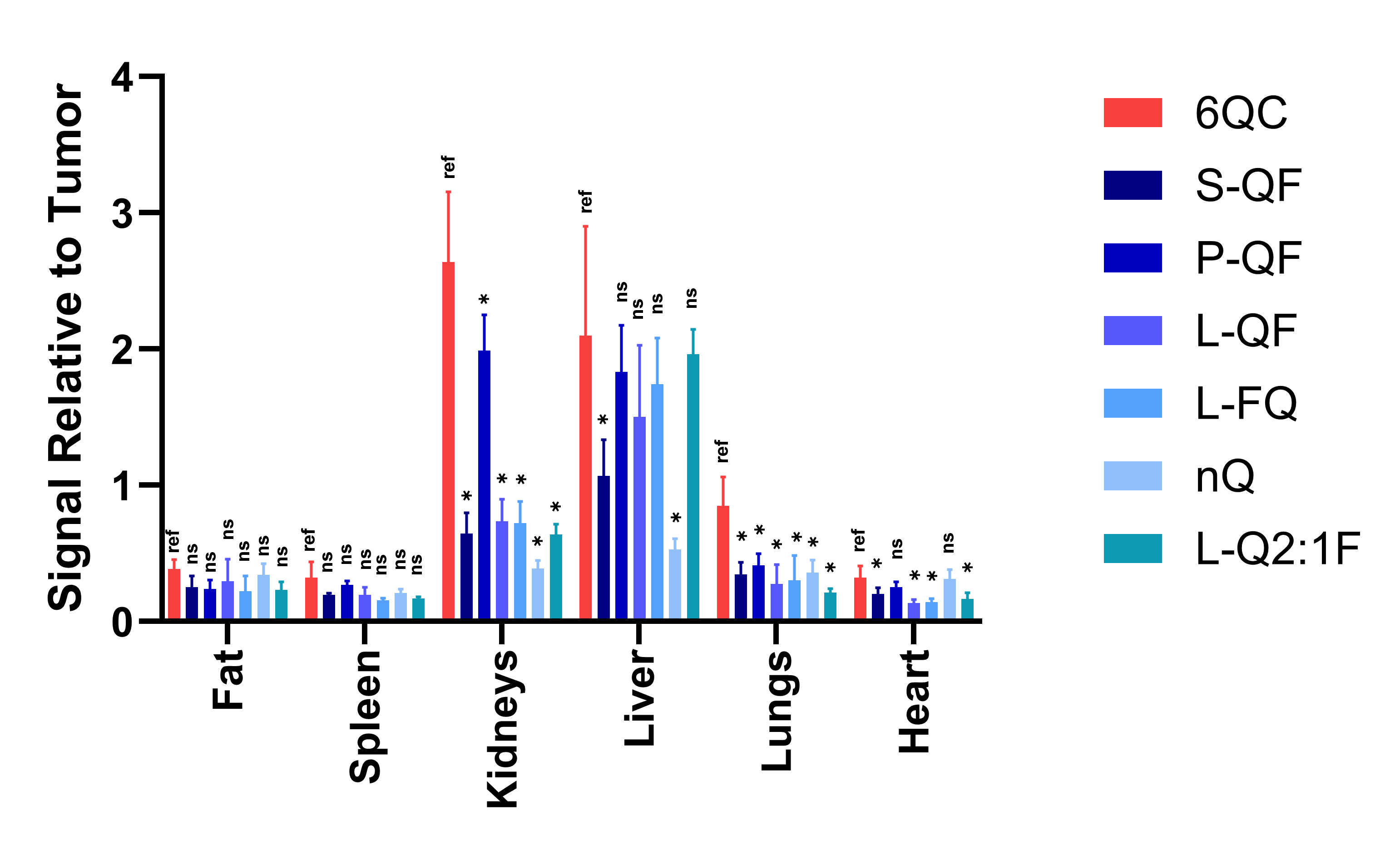

**Figure S1.** *Ex vivo* imaging of tumors and selected organs. Bar graph showing fluorescence distribution in selected organs. Images were acquired using small animal imaging system Pearl Trilogy and analyzed using Li-Cor Image Studio software. Statistics were calculated via Brown-Forsythe and Welch ANOVA test using GraphPad 10.2.0 (392), asterisk represents statistical significance (*p* $<0.05$), ns – not significant. *N:* 4 to 10 mice, 8 to 20 tumors per condition.

**Table S1.** Characterization of copolymer precursors: *M*_w_ – weight-average molecular weight; *M*_n_ – number-average molecular weight; *Ð* – polydispersity; TT content – molar percentage of the reactive thiazolidine-2-thione (TT) groups on the copolymer precursor.

| Precursor | *M*_w_  [kg.mol^-1^] | *M*_n_  [kg.mol^-1^] | *Ð* | TT content  [mol%] |
| --- | --- | --- | --- | --- |
| P1 | 64 | 54 | 1.19 | 12.1 |
| P2 | 66 | 57 | 1.16 | 13.3 |
| P3 | 65 | 54 | 1.20 | 11.6 |

**Table S2.** Compositions and characteristics of evaluated polymeric probes: *M*_w’_ – calculated as *M*_w_ of the copolymer precursor increased by the sum of apparent molecular weights of all conjugated ligands; F – weight percentage of the fluorophore moiety on the polymeric probe, as determined by amino acid analysis after polymer hydrolysis; No. of F – average number of fluorophore moieties per unit of polymer; Q – weight percentage of the quencher moiety on the polymeric probe, as determined by amino acid analysis after polymer hydrolysis; No. of Q – average number of quencher moieties per unit of polymer; Q:F Ratio – ratio between No. of Q and No. of F.

| Probe | Linker | Architecture | *M*_w’_  [kg.mol^-1^] | F  [wt%] | No. of F | Q  [wt%] | No. of Q | Q:F Ratio |
| --- | --- | --- | --- | --- | --- | --- | --- | --- |
| S-QF | PEG-3 | QF | 75 | 6.3 | 1.9 | - | - | - |
| S-Q1:1F | PEG-3 | Q 1:1 F | 78 | 5.1 | 2.8 | 4.5 | 3.5 | 1.3 |
| S-Q3:1F | PEG-3 | Q 3:1 F | 82 | 5.0 | 2.9 | 9.4 | 7.8 | 2.7 |
| P-QF | polyPro | QF | 73 | 9.9 | 2.1 | - | - | - |
| L-QF | PEG-12 | QF | 72 | 8.2 | 2.2 | - | - | - |
| L-FQ | PEG-12 | FQ | 67 | 7.7 | 1.8 | - | - | - |
| L-Q1:1F | PEG-12 | Q 1:1 F | 71 | 5.2 | 1.9 | 2.1 | 1.5 | 0.8 |
| L-Q2:1F | PEG-12 | Q 2:1 F | 75 | 6.1 | 2.4 | 6.2 | 4.7 | 1.9 |
| nQ | - | F | 67 | 2.7 | 2.3 | - | - | - |

**Table S3.** Evaluated probes and their *in vitro* characteristics (Initial cleavage rate rel. to 6QC and Signal-to-background ratio): *k*_rel_ – initial cleavage rate, relative to 6QC. Initial cleavage rate was evaluated as slope of the regression curve of the initial linear phase of the curve. *k*_rel_ was obtained as ratio between initial cleavage rate of the specified probe and 6QC; SBR - Signal-to-background ratio was evaluated as ratio between absorbance of the positive sample at 90 min and mean absorbance of negative samples at 90 min. All experiments were performed in triplicates.

| Probe | Linker | Architecture | *k*_rel_  [10 μM] | *k*_rel_  [2.5 μM] | SBR  [10 μM] | SBR  [2.5 μM] |
| --- | --- | --- | --- | --- | --- | --- |
| S-QF | PEG-4 | QF | 0.32 ± 0.01 | 0.39 ± 0.01 | 4.0 ± 0.3 | 5.0 ± 0.1 |
| S-Q1:1F | PEG-4 | Q1:1F | 0.32 ± 0.13 | 0.53 ± 0.09 | 1.3 ± 0.1 | 1.5 ± 0.1 |
| S-Q3:1F | PEG-4 | Q3:1F | 0.29 ± 0.01 | 0.52 ± 0.03 | 3.5 ± 0.1 | 4.1 ± 0.3 |
| P-QF | polyPro | QF | 0.51 ± 0.01 | 0.90 ± 0.03 | 15.2 ± 0.4 | 17.8 ± 0.2 |
| L-QF | PEG-12 | QF | 0.49 ± 0.01 | 1.24 ± 0.07 | 24.5 ± 0.7 | 33.7 ± 0.5 |
| L-FQ | PEG-12 | FQ | 0.45 ± 0.03 | 1.04 ± 0.09 | 29.2 ± 0.8 | 36.6 ± 1.5 |
| L-Q1:1F | PEG-12 | Q1:1F | 0.86 ± 0.05 | 2.18 ± 0.14 | 1.0 ± 0.1 | 1.3 ± 0.1 |
| L-Q2:1F | PEG-12 | Q2:1F | 0.77 ± 0.03 | 2.20 ± 0.07 | 3.0 ± 0.1 | 3.7 ± 0.1 |
| 6QC | - | - | 1 | 1 | 9.6 ± 0.5 | 9.9 ± 0.7 |

| Probe | Linker | Architecture | TBR  (2h) | TBR  (8h) | TBR  (24h) | TBR  (splay) |
| --- | --- | --- | --- | --- | --- | --- |
| S-QF | PEG-4 | QF | 2.5 ± 0.5 | 2.5 ± 0.5 | 2.7 ± 0.6 | 5.1 ± 1.9 |
| P-QF | polyPro | QF | 2.8 ± 0.6 | 3.1 ± 0.7 | 2.9 ± 0.4 | 4.7 ± 1.2 |
| L-QF | PEG-12 | QF | 3.0 ± 0.7 | 3.0 ± 0.9 | 3.1 ± 0.8 | 3.9 ± 1.5 |
| L-FQ | PEG-12 | FQ | 3.7 ± 1.0 | 3.6 ± 1.1 | 3.2 ± 0.3 | 3.4 ± 1.0 |
| L-Q2:1F | PEG-12 | Q2:1F | 2.9 ± 0.6 | 3.4 ± 0.4 | 3.9 ± 0.9 | 5.1 ± 1.3 |
| nQ | - | - | 2.1 ± 0.5 | 2.5 ± 0.5 | 3.0 ± 0.8 | 3.7 ± 1.4 |
| 6QC | - | - | 2.0 ± 0.4 | 2.1 ± 0.5 | 2.3 ± 0.5 | 2.3 ± 0.4 |

**Table S4.** Evaluated probes and their *in vivo* characteristics: TBR – tumor-to-background ratio. All values are show as mean ± standard deviation. Data are calculated from *N* = 4-10 mice (8-20 tumors) over multiple experiments.

### Methods

#### General Synthetic Methods

Unless stated otherwise all reactions were performed at ambient conditions. All solvents and reagents were used without further purification. All amino acids have L stereochemistry, standard Fmoc protected amino acids were used. To validate compound purity and identity (m/z) an analytical LC or LC-MS was used for all final compounds and key intermediates. The LC system used was HPLC (Jasco Inc.) equipped with a Reprosil 100 C18, 5 μm, 250 mm × 4 mm column. The LC-MS systems used was an Agilent 1200 HPLC with an Agilent Zorbax SB-C18 column (1.8 μm, 2.1 x 50 mm) using an Agilent 6125B Single Quad Mass Spectrometer or an Agilent 1100 Series HPLC with a Luna 4251-E0 C18 column (3 μm, 4.6 x 150 mm) coupled to a PE SCIEX API 3000 mass spectrometer. The wavelengths monitored were 215, 254 nm. All low-molecular weight compounds were purified using a CombiFlash 200 Rfi (Teledyne Isco) with a 4 or 12 g reverse phase C18 RediSep Rf Gold column or a preparatory Agilent 1260 Infinity HPLC system with a Luna Omega 5 μm C18 100A LC column 250 x 10 mm, which also monitored at wavelengths of 215, 254 nm. All gradient programs used a reverse phase gradient of 5-95% MeCN (0.1% TFA) in H_2_O (0.1% TFA) over 20-35 min unless otherwise stated. ESI MS and ESI HR-MS spectra were recorded on an Agilent 5975C MSD Quadrupole or LTQ Orbitrap XL (ThermoFisher Scientific).

**Materials**

1-amino-propan-2-ol, methacryloyl anhydride, methacryloyl chloride, thiazolidin-2-thione, β-alanine, *N*-ethyl-*N*′-(3-dimethylaminopropyl)carbodiimide hydrochloride (EDC.HCl), 2,5-di-*tert*-butylhydroquinone, carbon disulfide, ethanethiol, sodium hydride (60% dispersion in mineral oil), *N*,*N*-diisopropylethylamine (DIPEA), 4-(dimethylamino)pyridine (DMAP), *tert*-butanol (*t*-BuOH), *N*,*N*-dimethylacetamide (DMAA), dichloromethane (DCM), dimethyl sulfoxide (DMSO), 2,2′-azobisisobutyronitrile (AIBN) were from Merck. Initiator 2,2′-azobis(4-methoxy-2,4-dimethylvaleronitrile) (V-70) was obtained from FUJIFILM Wako Chemicals Europe GmbH (Germany). Sephadex LH-20 and Sephadex PD-10 columns was obtained from AP Czech Ltd. Sulfo-Cy5 was obtained from Lumiprobe Corp (USA). HBTU, HCTU and HATU were obtained from Peptides International Inc (USA). Trifluoroacetic acid (TFA) was obtained from Chem-Impex International (USA). 1,1,1,3,3,3-Hexafluoro-2-propanol (HFIP) was obtained from AA Blocks LLC (USA). 2,4,6-collidine was obtained from Alfa Aesar (USA). 2-Chlorotrityl chloride resin was obtained from AAPPTec (USA).

**Automatic Solid Phase Peptide Synthesis (SPPS)**

Peptides were synthesized using a Syro II (Biotage) fully automated parallel peptide synthesizer with standard reactor block. Resin used was 2-chlorotrityl chloride resin (1.05 mmol/g loading, 100-200 mesh) (AAPPTec, Cat. # RTZ001).

**Fmoc deprotection**: A 20% solution of piperidine in DMF was added to the reaction vessel for 5 min with agitation and drained. This was repeated once more.

**Resin washing**: Washing steps were completed between all coupling and deprotections. DMF was added to the reaction vessel, agitated, and then drained a total of three times.

**Amide bond couplings**: (2-(1Hbenzotriazol-1-yl)-1,1,3,3-tetramethyluronium hexafluorophosphate (HBTU), 2,4,6-collidine, and amino acids were pre-made into solutions using minimal DMF. 5 equivalents of HBTU, 10 equivalents of 2,4,6-collidine, and 5 equivalents of amino acid were added to the reaction vessel and incubated for 40 min with agitation. The mixture was then drained, and the coupling repeated for 40 min and then drained.

**Manual Solid Phase Peptide Synthesis (SPPS)**

2-Chlorotrityl chloride resin (256 mg, 1.5 mmol/g) was swelled in DCM (15 mL) for 15 min. Next, the resin was washed with DCM (2x, 15 mL) solution of FmocHN-hexane-NH_2_ (400 mg, 1.2 mmol) and DIPEA (410 μL, 2.3 mmol) in DMF (7 mL) was added. This reaction mixture was rocked overnight. Then, coupling solution was discarded and the resin was washed with DMF (3x, 15 mL), DCM (3x, 15 mL) and DMF (3x, 15 mL). Unbound beads were capped with MeOH (15 mL) for 50 min. MeOH was then discarded and the resin was washed again with DMF (3x, 15 mL), DCM (3x, 15 mL) and DMF (3x, 15 mL). Fmoc protecting group was removed with 20% piperidine in DMF (2x, 10 mL, 10 min) and the resin was washed with DMF (3x, 15 mL), DCM (3x, 15 mL) and DMF (3x, 15 mL). Next, coupling solution of Fmoc-Lys(Boc)-OH or Fmoc-Lys(Trt)-OH (1.2 mmol), HBTU (450 mg, 1.2 mmol) and 2,4,6-collidine (310 μL, 2.4 mmol) in DMF (10 mL) was added and the mixture was rocked for 1 h. The solution was then discarded, and the resin was washed with DMF (3x, 15 mL), DCM (3x, 15 mL) and DMF (3x, 15 mL).

The deprotection/coupling procedure was then repeated to extend the peptide chain to yield desired compounds. Acids and amino acids employed were: Fmoc-Phe-OH, Fmoc-Phg-OH, Fmoc-Ahx-OH, Boc-PEG-4-COOH. Modifications were introduced for coupling of Fmoc-Phg-OH to limit racemization. In this case, HBTU was replaced with HATU and Fmoc-Phg-peptide was deprotected using 20% 2-Methylpiperidine in DMF (2x, 10 mL, 10 min).

**Cleavage of Peptides from the Resin**

The resin was washed with DCM (5x, 15 mL) and the peptide was cleaved from the resin using DCM:HFIP (3:1, 10 mL). After 2 hours, the liquid was collected, and the resin was washed with DCM:HFIP (3:1, 2x, 5 mL). Volatiles were then evaporated, and the solid residue was resuspended in 0.1% TFA in H_2_O:MeCN (1:1, 30 mL) and freeze-dried to yield crude peptide. The peptides were used without further purification if not stated otherwise.

#### Synthesis of Target Linker-Modified Probe Derivatives

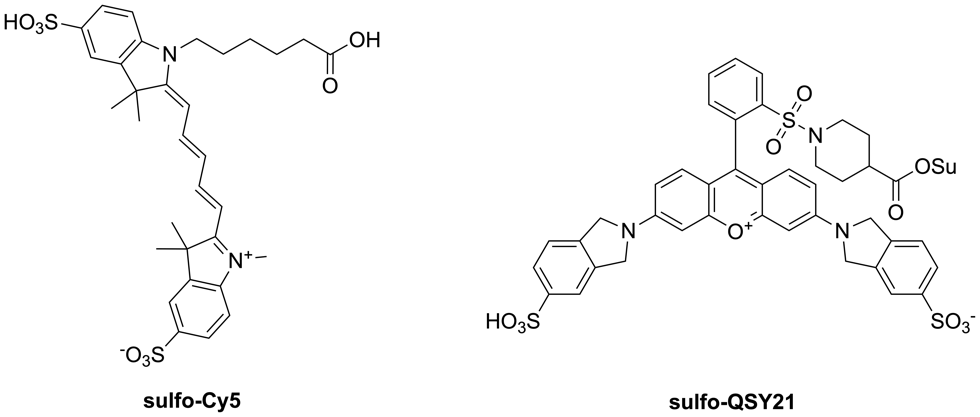

**Scheme S1: Structures of utilized dyes.** Structure of the fluorophore sulfo-Cy5 (left) and the quencher sulfo-QSY21 (right).

**Synthesis of S-QF Precursor 3:**

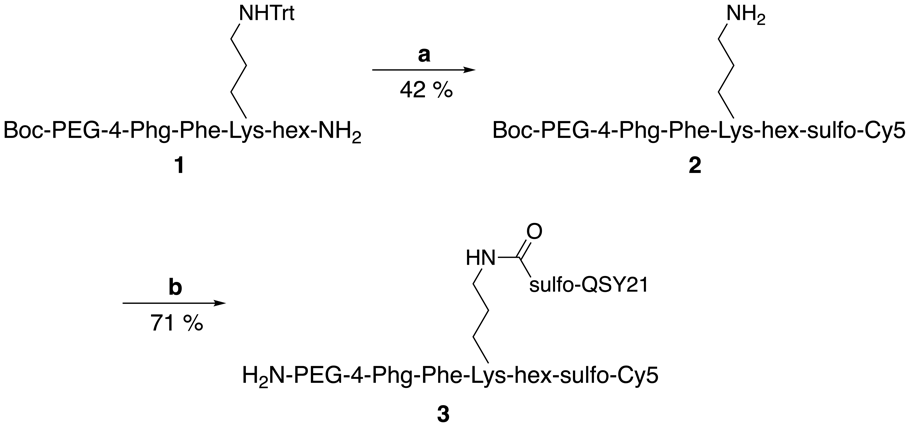

**Scheme S2: Synthesis of S-QF precursor: a)**; i) sulfo-Cy5-COOSu, DIPEA, DMF; ii) 1% TFA, DCM; **b)**; i) sulfo-QSY21-COOSu, DIPEA, DMF; ii) TFA, DCM.

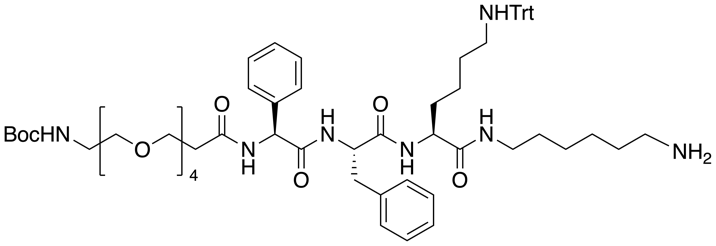

**Boc-PEG-4-Phg-Phe-Lys(Trt)-hex-NH_2_ (1):** Compound **1** was prepared according to general procedure for manual SPPS and cleavage of peptides from the resin. Boc-PEG-4-Phg-Phe-Lys(Trt)-hex-NH_2_ was isolated crude as white powder (273 mg, 53 %). 25.0 mg of the crude compound were then purified using (50 g C18, gradient 20-50% MeCN in H_2_O+0.1% TFA) to obtain 19.3 mg (77 %) of pure compound **1** as a white powder.

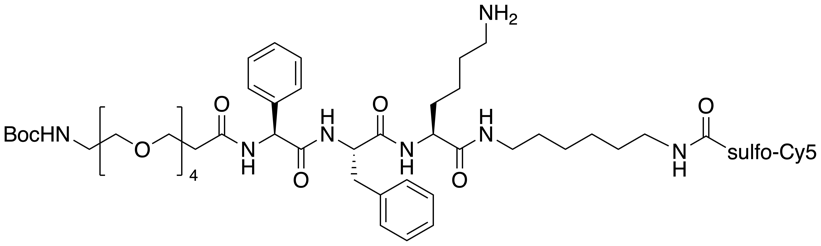

**Boc-PEG-4-Phg-Phe-Lys(H)-hex-sulfo-Cy5 (2):** Peptide **1** (6.5 mg, 4.9 μmol) was dissolved in DMF (1 mL) and DIPEA (4 μL, 25 μmol) was added to the mixture. Sulfo-Cyanine5 NHS ester (4.5 mg, 5.9 μmol) was then added and the reaction mixture was stirred at room temperature (rt). The reaction was followed by HPLC-MS until full consumption of starting materials. Then, volatiles were evaporated, and the residue was redissolved in 1% TFA in DCM (1 mL). The reaction was followed by HPLC-MS until full consumption of the Trt protected intermediate. Then, volatiles were evaporated again, and the residue was purified by flash chromatography (50 g C18, gradient 20-50% MeCN in H_2_O+0.1% TFA) to obtain compound **2** as blue powder (3.3 mg, 42 %).

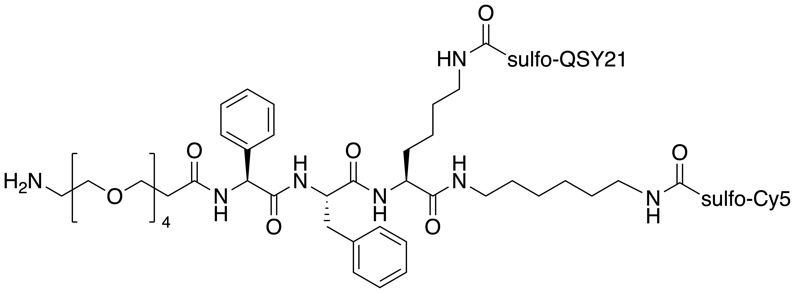

**H-PEG-4-Phg-Phe-Lys(sulfo-QSY21)-hex-sulfo-Cy5 (3):** Compound **2** (3.3 mg, 2.0 μmol) was dissolved in DMF (0.5 mL) and DIPEA (2 μL, 10 μmol) was added to the mixture. Sulfo-QSY21 NHS ester (2.8 mg, 3.0 μmol) was then added and the reaction mixture was stirred at rt. The reaction was followed by HPLC-MS until full consumption of starting materials. Then, volatiles were evaporated, and the residue was redissolved in DCM (0.5 mL) and TFA (0.5 mL). The reaction mixture was stirred for 15 min. Then, volatiles were evaporated again, and the residue was purified by flash chromatography (50 g C18, gradient 20-50% MeCN in H_2_O+0.1% TFA) to obtain compound **3** as blue powder (3.3 mg, 71 %). ESI MS: 1108.9 ([M $-$ 2H]^2-^). HR ESI MS: Calculated for C_113_H_132_O_25_N_12_S_5_ 1108.40205. Found 1108.40224.

**Synthesis of S-Qx:1F Precursor 5:**

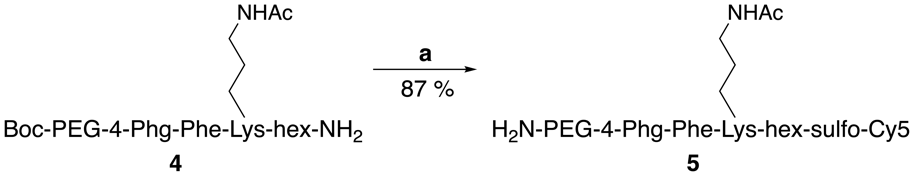

**Scheme S3: Synthesis of S-Qx:1F precursor: a)**; i) sulfo-Cy5-COOSu, DIPEA, DMF; ii) TFA, DCM.

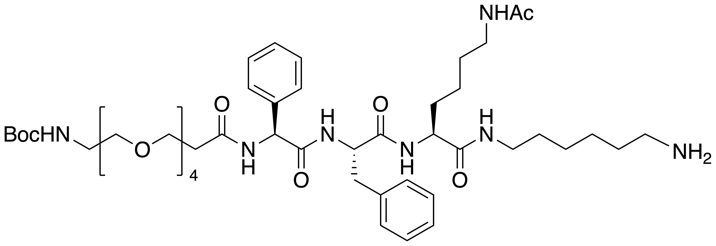

**Boc-PEG-4-Phg-Phe-Lys(Ac)-hex-NH_2_ (4):** This compound was prepared according to general procedure for manual SPPS and cleavage of peptides from the resin. Boc-PEG-4-Phg-Phe-Lys(Ac)-hex-NH_2_ was isolated crude as white powder (197 mg, 45 %). 12.0 mg of the crude compound were then purified using (50 g C18, gradient 20-50% MeCN in H_2_O+0.1% TFA) to obtain 9.3 mg (78 %) of pure compound **4** as a white powder.

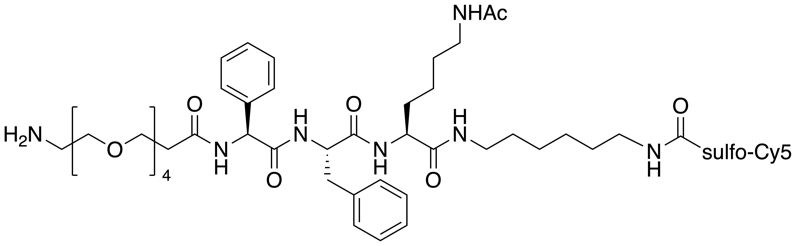

**H-PEG-4-Phg-Phe-Lys(Ac)-hex-sulfo-Cy5 (5):** Peptide **4** (9.3 mg, 8.2 μmol) was dissolved in DMF (1 mL) and DIPEA (7 μL, 41 μmol) was added to the mixture. Sulfo-Cyanine5 NHS ester (7.5 mg, 9.8 μmol) was then added and the reaction mixture was stirred at rt. The reaction was followed by HPLC-MS until full consumption of starting materials. Then, volatiles were evaporated, and the residue was redissolved in DCM (0.5 mL) and TFA (0.5 mL). The reaction mixture was stirred for 15 min. Then, volatiles were evaporated again and the residue was purified by flash chromatography (50 g C18, gradient 20-40% MeCN in H_2_O+0.1% TFA) to obtain compound **5** as blue powder (11 mg, 87 %). ESI MS: 1438.7 ([M $+$ H]^+^). HR ESI MS: Calculated for C_74_H_105_O_16_N_9_S_2_ 719.85549. Found 719.85536.

**Synthesis of Q Precursor 7:**

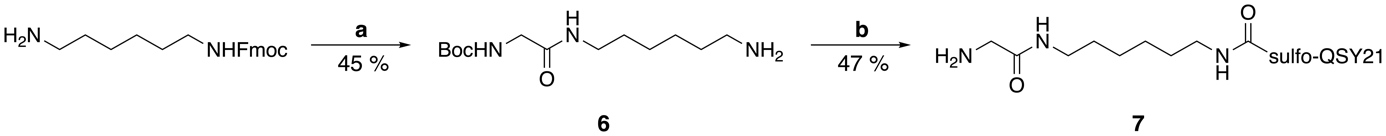

**Scheme S4: Synthesis of Q precursor: a)**; i) Boc-Gly-OH, HCTU, 2,4,6-collidine, DMF; ii) 20% piperidine, DMF; **b)**; i) sulfo-QSY21-COOSu, DIPEA, DMF; ii) TFA, DCM.

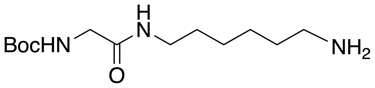

**Boc-Gly-hex-NH_2_ (6):** Boc-Gly-OH (187 mg, 1.1 mmol) and HCTU (440 mg, 1.1 mmol) were dissolved in DMF (5 mL). 2,4,6-Collidine (0.85 mL, 6.4 mmol) was added, and the reaction mixture was stirred for 5 min at rt. Next, *N*-Fmoc-1,6-diaminohexane (400 mg, 1.1 mmol) was added and the reaction was stirred overnight. Then, solvent was evaporated, DCM was added (20 mL), and the organic phase was washed with 10% KHSO_4_ (2x20 mL), distilled H_2_O (2x20 mL), sat. NaHCO_3_ (2x20 mL) and brine (10 mL). Then, volatiles were evaporated, and the residue was redissolved in 20% piperidine in DMF (5 mL). The reaction was followed by HPLC-MS until full consumption of the Fmoc-protected intermediate. Volatiles were evaporated and the residue was purified by flash chromatography (50 g C18, gradient 10-40% MeCN in H_2_O+0.1% TFA) to obtain compound **6** as white powder (187 mg, 45 %).

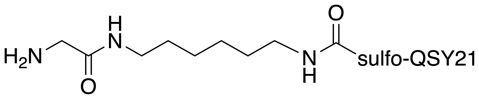

**H-Gly-hex-sulfo-QSY21 (7):** Compound **6** (6.0 mg, 22 μmol) was dissolved in DMF (1 mL) and DIPEA (57 μL, 329 μmol) was added to the mixture. QSY21 NHS ester (10 mg, 11 μmol) was then added and the reaction was stirred at rt. The reaction was followed by HPLC-MS until full consumption of starting materials. Then, volatiles were evaporated, and the residue was redissolved in DCM (0.5 mL) and TFA (0.5 mL). The reaction mixture was stirred for 15 min. Then, volatiles were evaporated again, and the product was purified by flash chromatography (50 g C18, gradient 20-60% MeCN in H_2_O+0.1% TFA) to obtain compound **7** as blue powder (5.2 mg, 47 %). ESI MS: 995.3 ([M $-$ H]^-^). HR ESI MS: Calculated for C_49_H_51_O_11_N_6_S_3_ 995.27834. Found 995.27899.

**Synthesis of P-QF Precursor 11:**

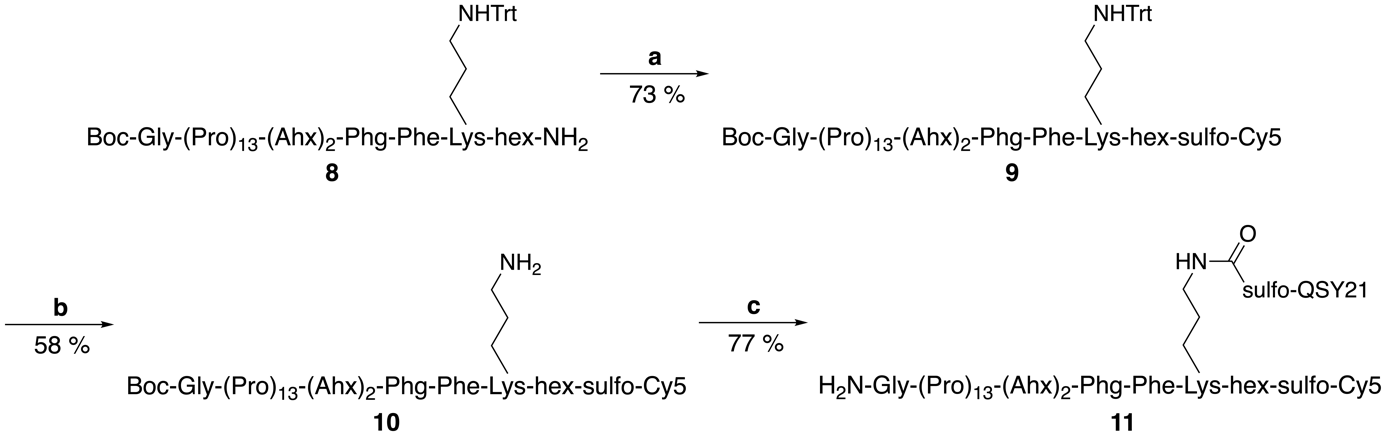

**Scheme S5: Synthesis of P-QF precursor: a)** sulfo-Cy5-COOSu, DIPEA, DMF; **b)** 1% TFA, DCM; **c)**; i) sulfo-QSY21-COOSu, DIPEA, DMF; ii) TFA, DCM.

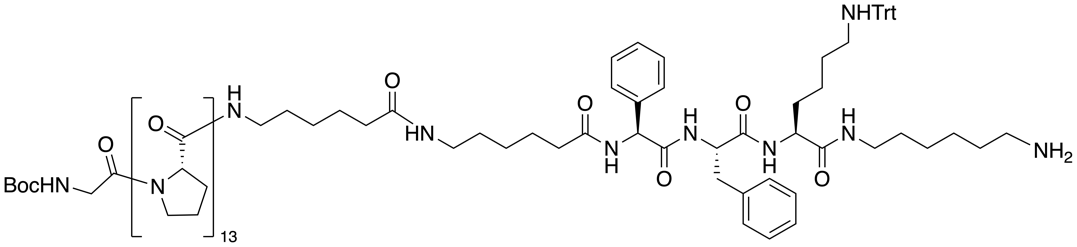

**Boc-Gly-Pro_13_-Ahx-Ahx-Phg-Phe-Lys(Trt)-hex-NH_2_ (8):** This compound was prepared according to general procedure for SPPS and cleavage of peptides from the resin. The Fmoc-Ahx-Ahx-Phg-Phe-Lys(Trt)-hex-resin fragment was prepared using manual SPPS and was then extended to final Boc-Gly-Pro_13_-Ahx-Ahx-Phg-Phe-Lys(Trt)-hex-NH_2_ using automatic SPPS. It was isolated crude as white powder (225 mg, 57 %). 10.0 mg of the crude compound were then purified using (50 g C18, gradient 20-50% MeCN in H_2_O+0.1% TFA) to obtain 8.1 mg (81 %) of pure compound **8** as a white powder.

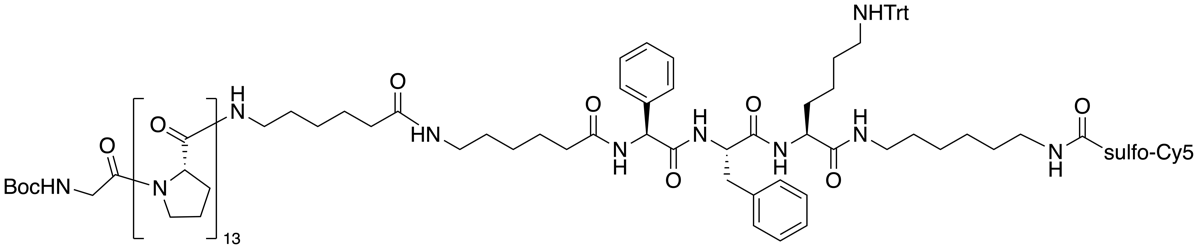

**Boc-Gly-Pro_13_-Ahx-Ahx-Phg-Phe-Lys(Trt)-hex-sulfo-Cy5 (9):** Peptide **8** (8.0 mg, 3.2 μmol) was dissolved in DMF (1 mL) and DIPEA (3 μL, 16 μmol) was added to the mixture. Sulfo-Cyanine5 NHS ester (4.7 mg, 6.4 μmol) was then added and the reaction mixture was stirred at rt. The reaction was followed by HPLC-MS until full consumption of starting materials. Then volatiles were evaporated, and the residue was purified by flash chromatography (50 g C18, gradient 20-60% MeCN in H_2_O+0.1% TFA) to obtain compound **9** as blue powder (7.1 mg, 73 %).

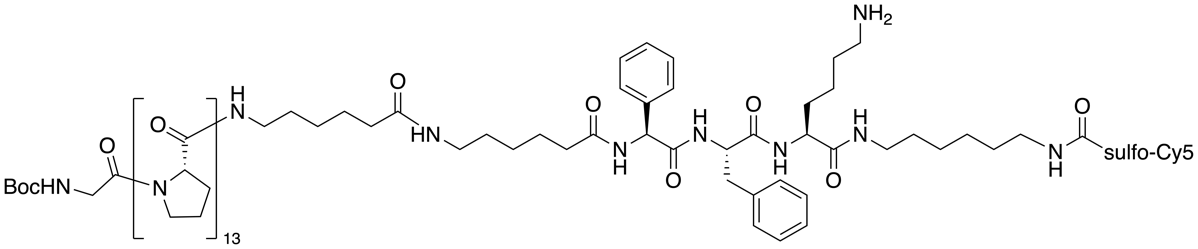

**Boc-Gly-Pro_13_-Ahx-Ahx-Phg-Phe-Lys(H)-hex-sulfo-Cy5 (10):** Compound **9** (7.1 mg, 2.3 μmol) was dissolved in 1% TFA in DCM (1 mL). The reaction was followed by HPLC-MS until full consumption of the Trt protected intermediate and then volatiles were evaporated. The residue was purified by flash chromatography (50 g C18, gradient 20-50% MeCN in H_2_O+0.1% TFA) to obtain compound **10** as blue powder (3.9 mg, 58 %).

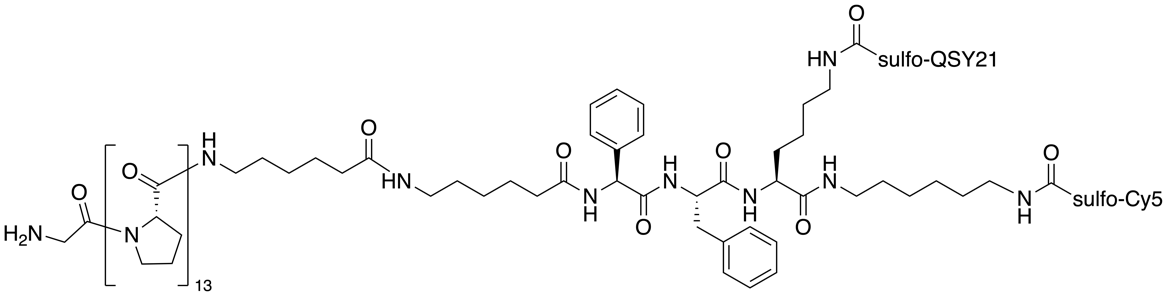

**H-Gly-Pro_13_-Ahx-Ahx-Phg-Phe-Lys(sulfo-QSY21)-hex-sulfo-Cy5 (11):** Compound **10** (3.9 mg, 1.3 μmol) was dissolved in DMF (0.5 mL) and DIPEA (2 μL, 10 μmol) was added to the mixture. Sulfo-QSY21 NHS ester (2.4 mg, 2.6 μmol) was then added and the reaction mixture was stirred at rt. The reaction was followed by HPLC-MS until full consumption of starting materials. Then, volatiles were evaporated, and the residue was redissolved in DCM (0.5 mL) and TFA (0.5 mL). The reaction mixture was stirred for 15 min Then, volatiles were evaporated again. The residue was purified by flash chromatography (50 g C18, gradient 20-60% MeCN in H_2_O+0.1% TFA) to obtain compound **11** as blue powder (3.6 mg, 77 %). Boc-protected precursor ESI MS: 1207.7 ([M $+$ 3H]^3+^).

**Synthesis of L-QF Precursor 16:**

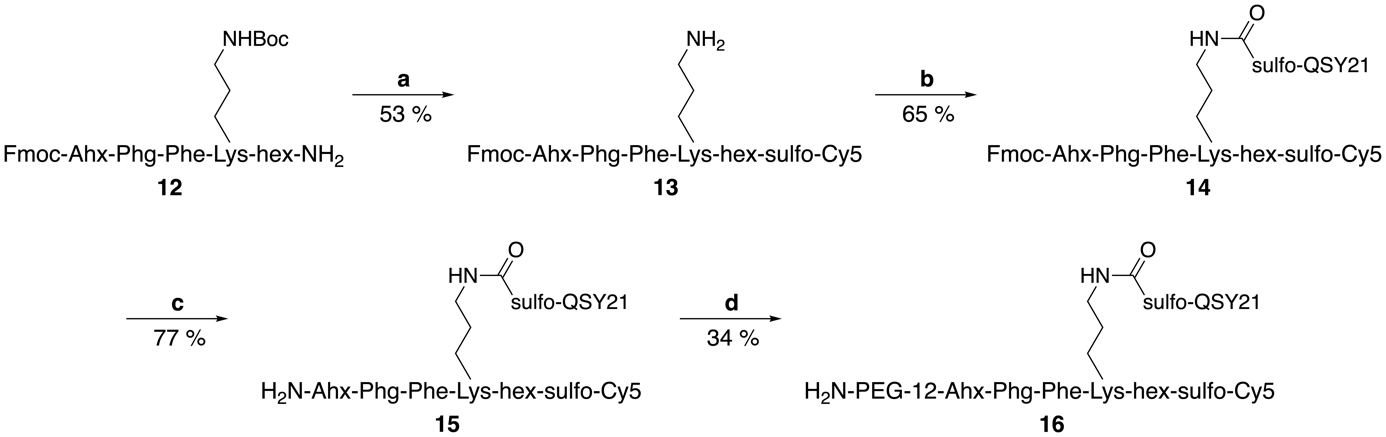

**Scheme S6: Synthesis of L-QF precursor: a)**; i) sulfo-Cy5-COOSu, DIPEA, DMF; ii) TFA, DCM; **b)** sulfo-QSY21-COOSu, DIPEA, DMF; **c)** 1% DBU, DMF; **d)**; i) Boc-PEG-12-CH_2_CH_2_COOH, HATU, 2,4,6-collidine, DMF; ii) TFA, DCM.

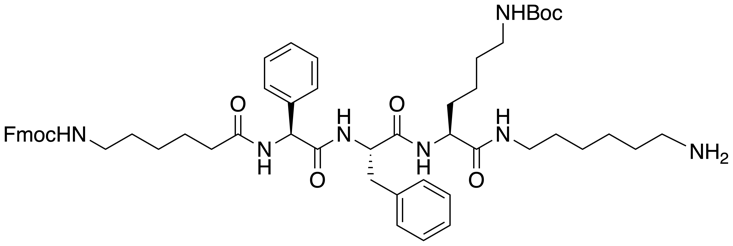

**Fmoc-Ahx-Phg-Phe-Lys(Boc)-hex-NH_2_ (12):** This compound was prepared according to general procedure for manual SPPS and cleavage of peptides from the resin. Fmoc-Ahx-Phg-Phe-Lys(Ac)-hex-NH_2_ was isolated as white powder (175 mg, 49 %).

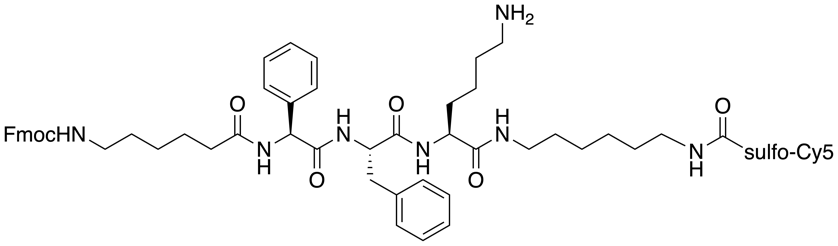

**Fmoc-Ahx-Phg-Phe-Lys(H)-hex-sulfo-Cy5 (13):** Peptide **12** (10 mg, 9.3 μmol) was dissolved in DMF (1 mL) and DIPEA (9 μL, 47 μmol) was added to the mixture. Sulfo-Cyanine5 NHS ester (8.7 mg, 12 μmol) was then added and the reaction mixture was stirred at rt. The reaction was followed by HPLC-MS until full consumption of starting materials. Next, volatiles were evaporated, and the residue was redissolved in DCM (0.5 mL) and TFA (0.5 mL). The reaction mixture was stirred for 30 min. Then, volatiles were evaporated again. The residue was purified by flash chromatography (50 g C18, gradient 20-60% MeCN in H_2_O+0.1% TFA) to obtain compound **13** as blue powder (8.0 mg, 53 %).

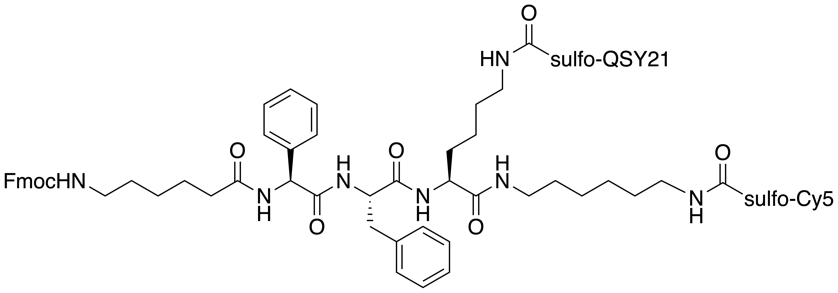

**Fmoc-Ahx-Phg-Phe-Lys(sulfo-QSY21)-hex-sulfo-Cy5 (14):** Compound **13** (8.0 mg, 5.0 μmol) was dissolved in DMF (1 mL) and DIPEA (9 μL, 50 μmol) was added to the mixture. Sulfo-QSY21 NHS ester (6.6 mg, 7.0 μmol) was then added and the reaction mixture was stirred at rt. The reaction was followed by HPLC-MS until full consumption of starting materials. Next, the product was purified by flash chromatography (50 g C18, gradient 20-50% MeCN in H_2_O+0.1% TFA) to obtain compound **14** as blue powder (7.5 mg, 65 %).

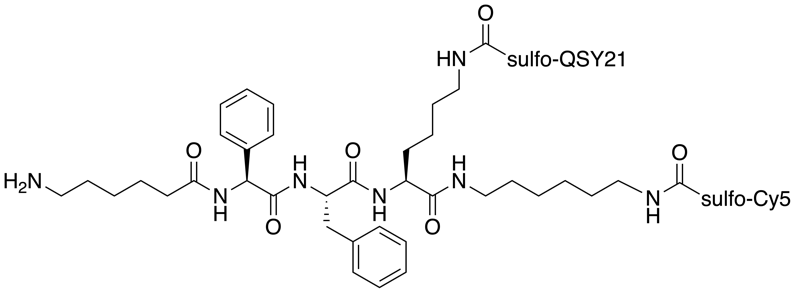

**H-Ahx-Phg-Phe-Lys(sulfo-QSY21)-hex-sulfo-Cy5 (15):** Compound **14** (7.5 mg, 3.2 μmol) was dissolved in 1% DBU in DMF (1 mL) and stirred at rt. Reaction was quenched after 15 min by addition of AcOH (20 μL). Next, the product was purified by flash chromatography (50 g C18, gradient 20-50% MeCN in H_2_O+0.1% TFA) to obtain compound **15** as blue powder (5.5 mg, 77 %).

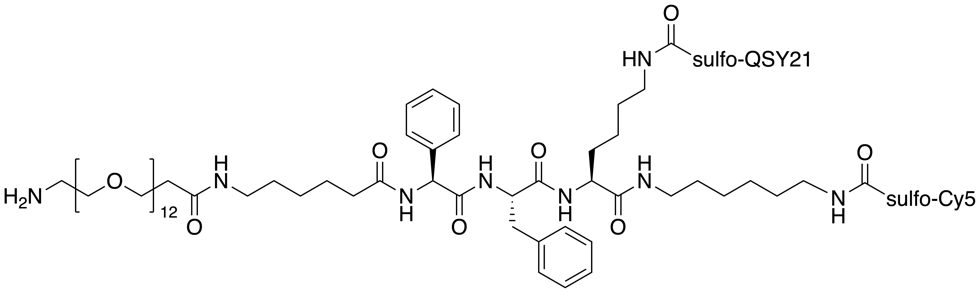

**H-PEG-12-OCH_2_CH_2_-Ahx-Phg-Phe-Lys(QSY21)-hex-sulfo-Cy5 (16) :**BocHN-PEG-12-OCH_2_CH_2_COOH (3.6 mg, 5.0 μmol) and HATU (1.8 mg, 4.8 μmol) were dissolved in DMF (1 mL). 2,4,6-Collidine (3.3 μL, 25 μmol) was added and the reaction mixture was stirred for 5 min at rt. Next, amine **15** (5.5 mg, 2.5 μmol) was added and the reaction was stirred for 3 hours. Then, volatiles were evaporated, and the residue was redissolved in DCM (0.5 mL) and TFA (0.5 mL). The reaction mixture was stirred for 15 min. Then, volatiles were evaporated again. The residue was purified by flash chromatography (50 g C18, gradient 20-60% MeCN in H_2_O+0.1% TFA) to obtain compound **16** as blue powder (2.4 mg, 34 %). ESI MS: 1341.1 ([M $-$ 2H]^2-^). HR ESI MS: Calculated for C_135_H_175_O_34_N_13_S_5_ 1341.04894. Found 1341.04940.

**Synthesis of L-FQ Precursor 20:**

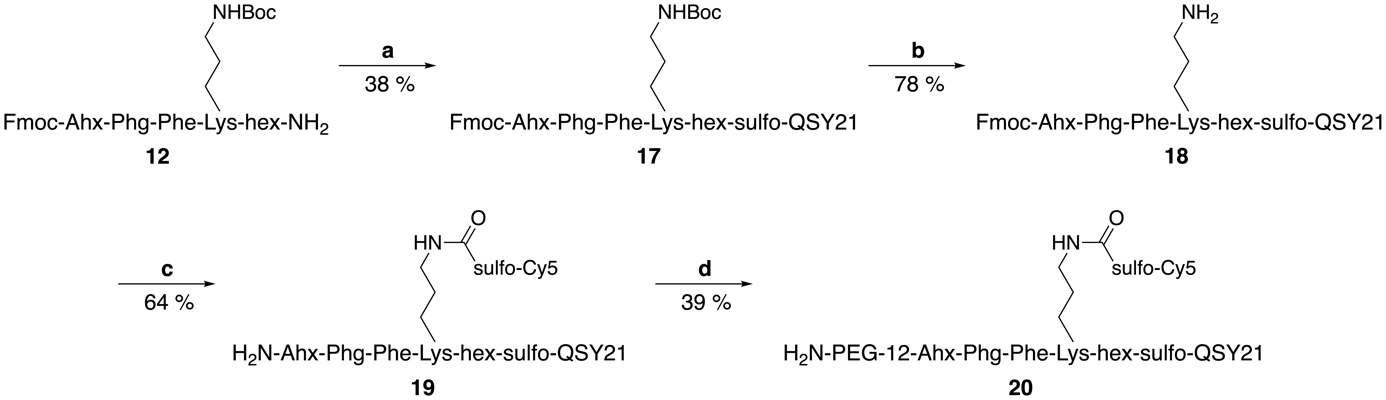

**Scheme S7: Synthesis of L-FQ precursor: a)** sulfo-QSY21-COOH, PyBOP, DIPEA, DMF; **b)** TFA, DCM; **c)**; i) sulfo-Cy5-COOSu, DIPEA, DMF; ii) 1% DBU; **d)**; i) Boc-PEG-12-CH_2_CH_2_COOH, HATU, 2,4,6-collidine, DMF; ii) TFA, DCM.

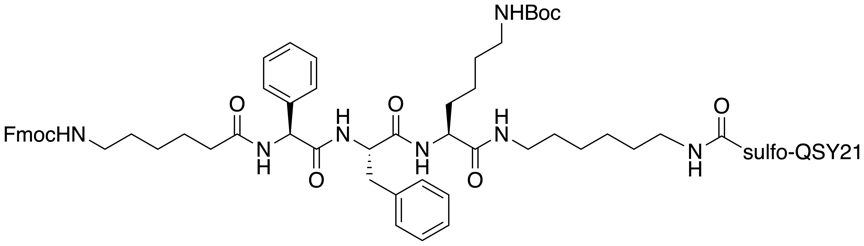

**Fmoc-Ahx-Phg-Phe-Lys(Boc)-hex-sulfo-QSY21 (17):** Peptide **12** (13 mg, 12 μmol), sulfo-QSY21 (10 mg, 12 μmol) and PyBOP (8.0 mg, 15 μmol) were dissolved in DMF (2 mL). Then, DIPEA (17 μL, 95 μmol) was added to the mixture and the reaction mixture was stirred at rt. The reaction was followed by HPLC-MS until full consumption of starting materials. Next, volatiles were evaporated and, the residue was purified by flash chromatography (50 g C18, gradient 20-60% MeCN in H_2_O+0.1% TFA) to obtain compound **17** as blue powder (8.0 mg, 38 %).

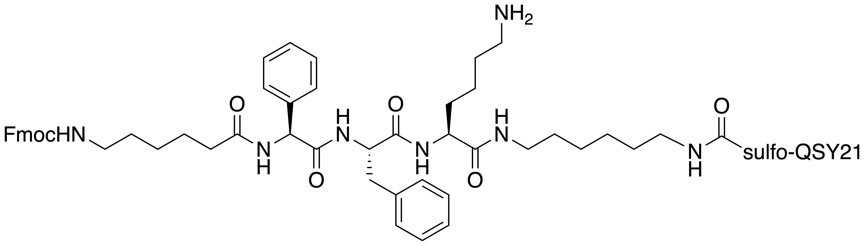

**Fmoc-Ahx-Phg-Phe-Lys(H)-hex-sulfo-QSY21 (18):** Compound **17** (23 mg, 13 μmol) was dissolved in DCM (0.5 mL) and TFA (0.5 mL). The reaction mixture was stirred for 30 min. Then, volatiles were evaporated, and the residue was purified by flash chromatography (50 g C18, gradient 20-50% MeCN in H_2_O+0.1% TFA) to obtain compound **18** as blue powder (18 mg, 78 %).

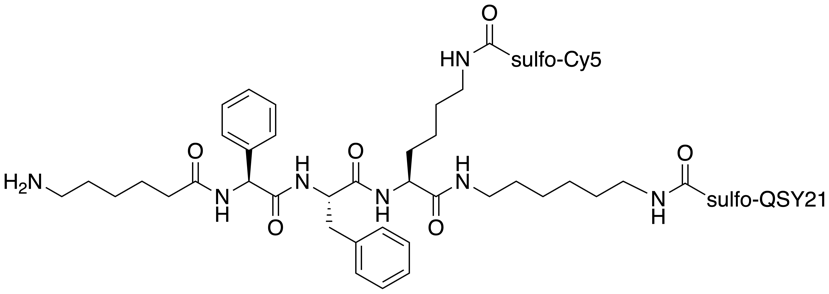

**H-Ahx-Phg-Phe-Lys(Cy5)-hex-sulfo-QSY21 (19):** Compound **18** (18 mg, 10 μmol) was dissolved in DMF (2 mL) and DIPEA (26 μL, 150 μmol) was added to the mixture. Sulfo-Cyanine5 NHS ester (8.9 mg, 12 μmol) was then added and the reaction mixture was stirred at rt. The reaction was followed by HPLC-MS until full consumption of starting materials. Next, 2% solution of DBU in DMF (2 mL) was added and the reaction mixture was stirred for 15 min. Then, 150 μL of AcOH were added to quench the reaction. Then, volatiles were evaporated, and the residue was purified by flash chromatography (50 g C18, gradient 10-40% MeCN in H_2_O+0.1% TFA) to obtain compound **19** as blue powder (14 mg, 64 %).

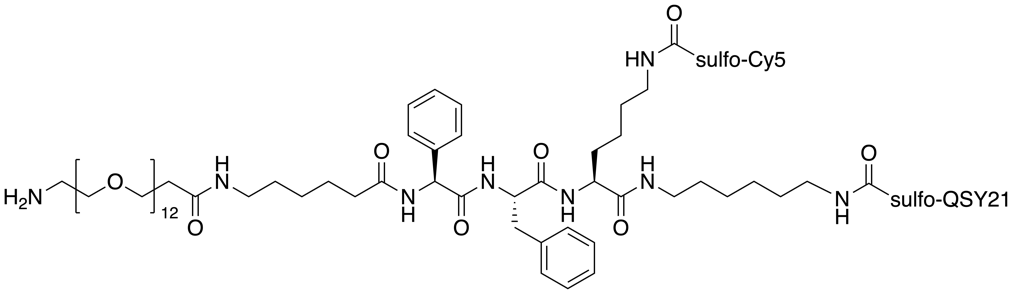

**H-PEG-12-OCH_2_CH_2_-Ahx-Phg-Phe-Lys(Cy5)-hex-sulfo-QSY21 (20):** BocHN-PEG-12-OCH_2_CH_2_COOH (9.1 mg, 13 μmol) and HATU (4.4 mg, 11 μmol) were dissolved in DMF (2 mL). 2,4,6-Collidine (13 μL, 95 μmol) was added and the reaction mixture was stirred for 5 min at rt. Next, amine **19** (14 mg, 6.4 μmol) was added and the reaction was stirred for 3 hours. Then, volatiles were evaporated, and the residue was redissolved in DCM (0.5 mL) and TFA (0.5 mL). The reaction mixture was stirred for 15 min. Then, volatiles were evaporated again, and the residue was purified by flash chromatography (50 g C18, gradient 15-50% MeCN in H_2_O+0.1% TFA) to obtain compound **20** as blue powder (7.0 mg, 39 %). ESI MS: 1341.6 ([M $-$ 2H]^2-^). HR ESI MS: Calculated for C_135_H_175_O_34_N_13_S_5_ 1341.04894. Found 1341.04955.

**Synthesis of L-Qx:1F Precursor 23:**

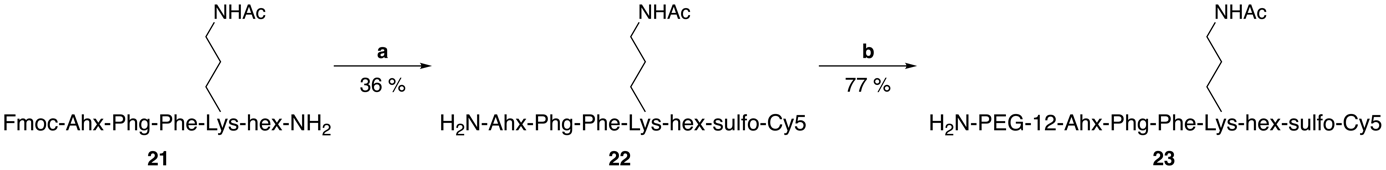

**Scheme S8: Synthesis of L-Qx:1F precursor: a)**; i) sulfo-Cy5-COOSu, DIPEA, DMF; ii) 1% DBU; **b)**; i) Boc-PEG-12-CH_2_CH_2_COOH, HATU, 2,4,6-collidine, DMF; ii) TFA, DCM.

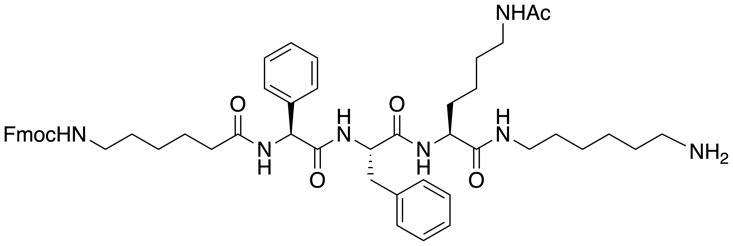

**Fmoc-Ahx-Phg-Phe-Lys(Ac)-hex-NH_2_ (21):** This compound was prepared according to general procedure for manual SPPS and cleavage of peptides from the resin. Fmoc-Ahx-Phg-Phe-Lys(Ac)-hex-NH_2_ was isolated as white powder (175 mg, 51 %).

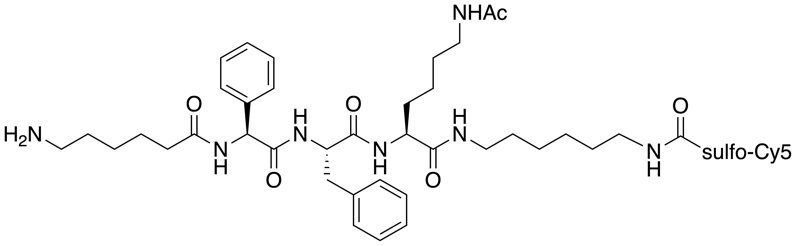

**H-Ahx-Phg-Phe-Lys(Ac)-hex-sulfo-Cy5 (22):** Peptide **21** (10 mg, 9.8 μmol) was dissolved in DMF (1 mL) and DIPEA (9 μL, 49 μmol) was added to the mixture. Sulfo-Cyanine5 NHS ester (9.2 mg, 12 μmol) was then added and the reaction mixture was stirred at rt. The reaction was followed by HPLC-MS until full consumption of starting materials. Next, DBU (10 uL) was added to achieve Fmoc deprotection and the reaction mixture was stirred for 15 min at rt. It was then quenched by addition of AcOH (20 μL) and purified by flash chromatography (50 g C18, gradient 20-40% MeCN in H_2_O+0.1% TFA) to obtain compound **22** as blue powder (5.0 mg, 36 %).

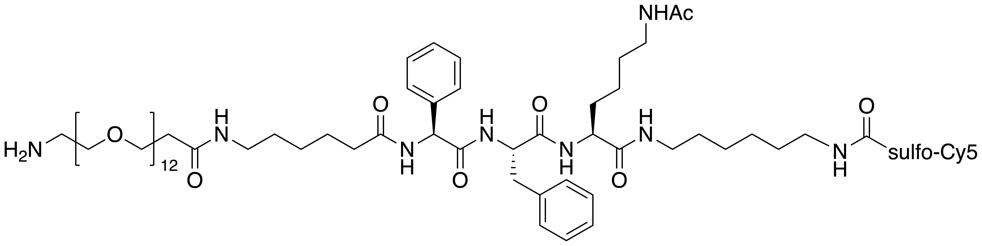

**H-PEG-12-OCH_2_CH_2_-Ahx-Phg-Phe-Lys(Ac)-hex-sulfo-Cy5 (23):** BocHN-PEG-12-OCH_2_CH_2_COOH (5.0 mg, 7.0 μmol) and HATU (2.4 mg, 6.3 μmol) were dissolved in DMF (1 mL). 2,4,6-Collidine (5.0 μL, 35 μmol) was added and the reaction mixture was stirred for 5 min at rt. Next, amine **22** (5.0 mg, 3.5 μmol) was added and the reaction was stirred for 1 hour. Then, volatiles were evaporated, and the residue was redissolved in DCM (0.5 mL) and TFA (0.5 mL). The reaction mixture was stirred for 15 min. Then, volatiles were evaporated again, and the residue was purified by flash chromatography (50 g C18, gradient 20-40% MeCN in H_2_O+0.1% TFA) to obtain compound **23** as blue powder (5.5 mg, 77 %). ESI MS: 951.0 ([M $-$ 2H]^2-^). HR ESI MS: Calculated for C_96_H_144_O_25_N_10_S_2_ 950.48782. Found 950.48743.

**Synthesis of nQ Precursor 25:**

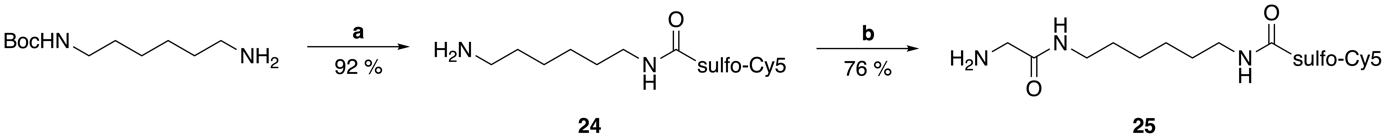

**Scheme S9: Synthesis of nQ precursor: a)**; i) sulfo-Cy5-COOSu, DIPEA, DMF; ii) TFA, DCM; **b)**; i) Boc-Gly-OSu, DIPEA, DMF; ii) TFA, DCM.

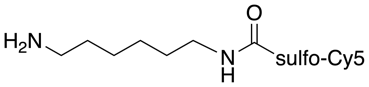

**H-hex-sulfo-Cy5 (24)**: *N*-Boc-1,6-diaminohexane (7.0 mg, 32 μmol) was dissolved in DMF (1 mL) and DIPEA (24 μL, 135 μmol) was added to the mixture. Sulfo-Cyanine5 NHS ester (20 mg, 27 μmol) was then added and the reaction was stirred at rt. The reaction was followed by HPLC-MS until full consumption of starting materials. Then, volatiles were evaporated, and the residue was redissolved in DCM (0.5 mL) and TFA (0.5 mL). The reaction mixture was stirred for 15 min. Then, volatiles were evaporated again, and the product was purified by flash chromatography (50 g C18, gradient 10-40% MeCN in H_2_O+0.1% TFA) to obtain compound **24** as blue powder (21 mg, 92 %).

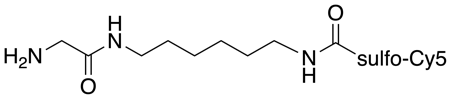

**H-Gly-hex-sulfo-Cy5 (25):** Compound **24** (16 mg, 19 μmol) was dissolved in DMF (1 mL) and DIPEA (16 μL, 93 μmol) was added to the mixture. Boc-Gly-OSu (6.1 mg, 22 μmol) was then added and the reaction was stirred at rt. The reaction was followed by HPLC-MS until full consumption of starting materials. Then, volatiles were evaporated, and the residue was redissolved in DCM (0.5 mL) and TFA (0.5 mL). The reaction mixture was stirred for 15 min. Next, volatiles were evaporated again, and the product was purified by flash chromatography (50 g C18, gradient 10-40% MeCN in H_2_O+0.1% TFA) to obtain compound **25** as blue powder (13 mg, 76 %). ESI MS: 796.3 ([M $-$ H]^-^). HR ESI MS: Calculated for C_40_H_54_O_8_N_5_S_2_ 796.34193. Found 796.34137.

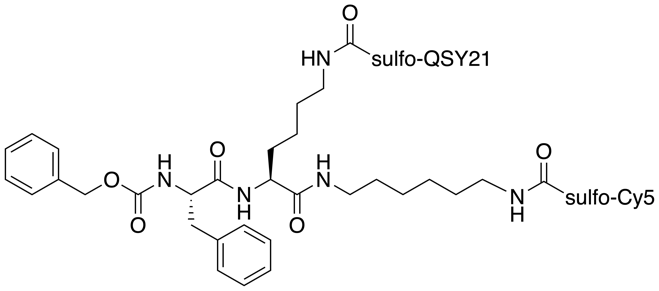

**6QC (26):** 6QC was synthesized according to a published procedure.^1^

#### HPLC Chromatograms of Target Linker-Modified Probe Derivatives

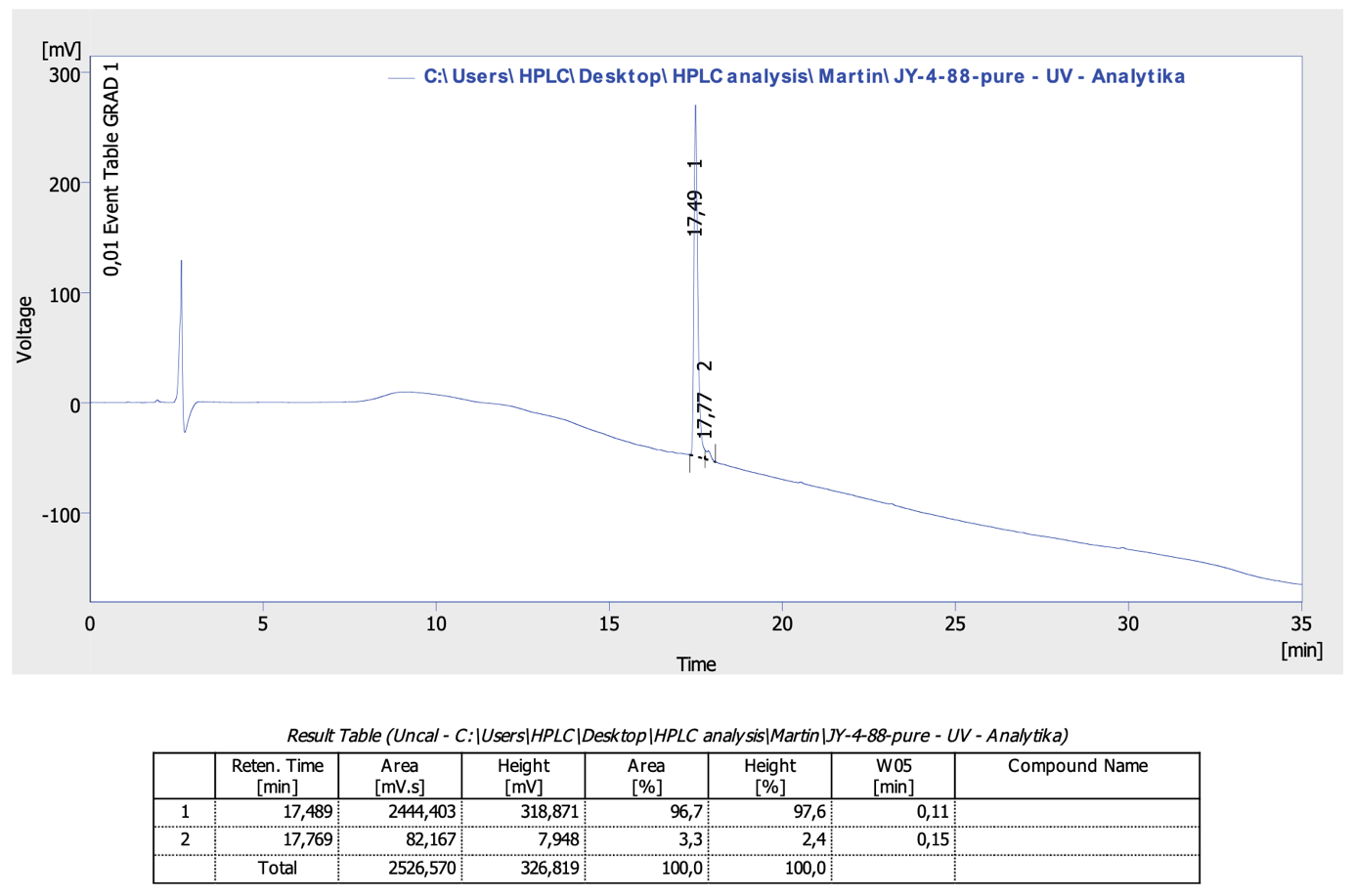

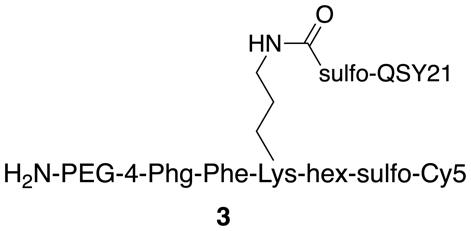

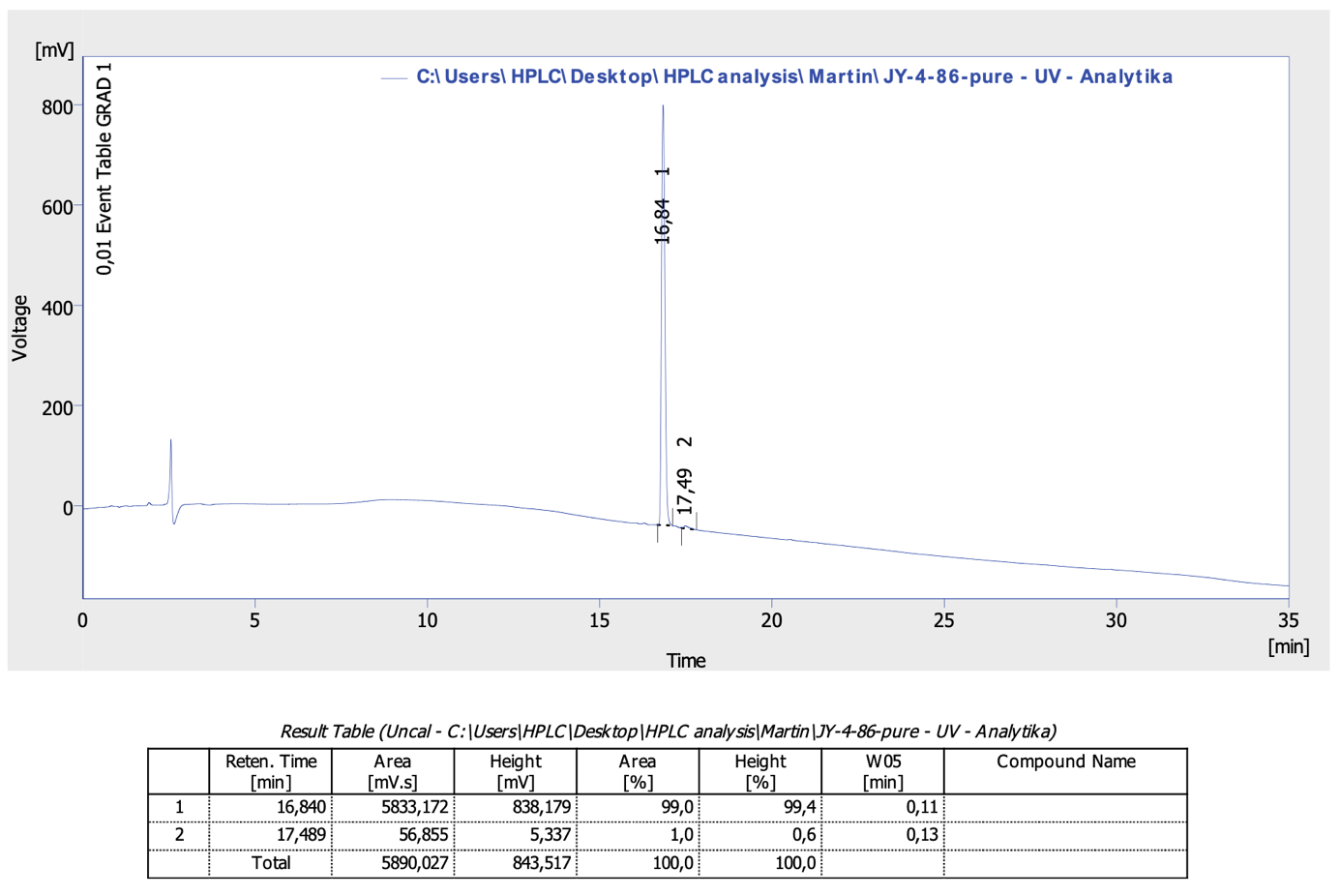

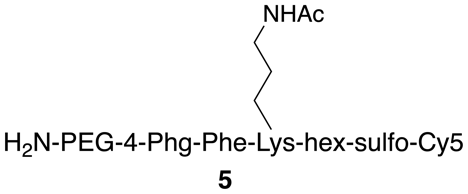

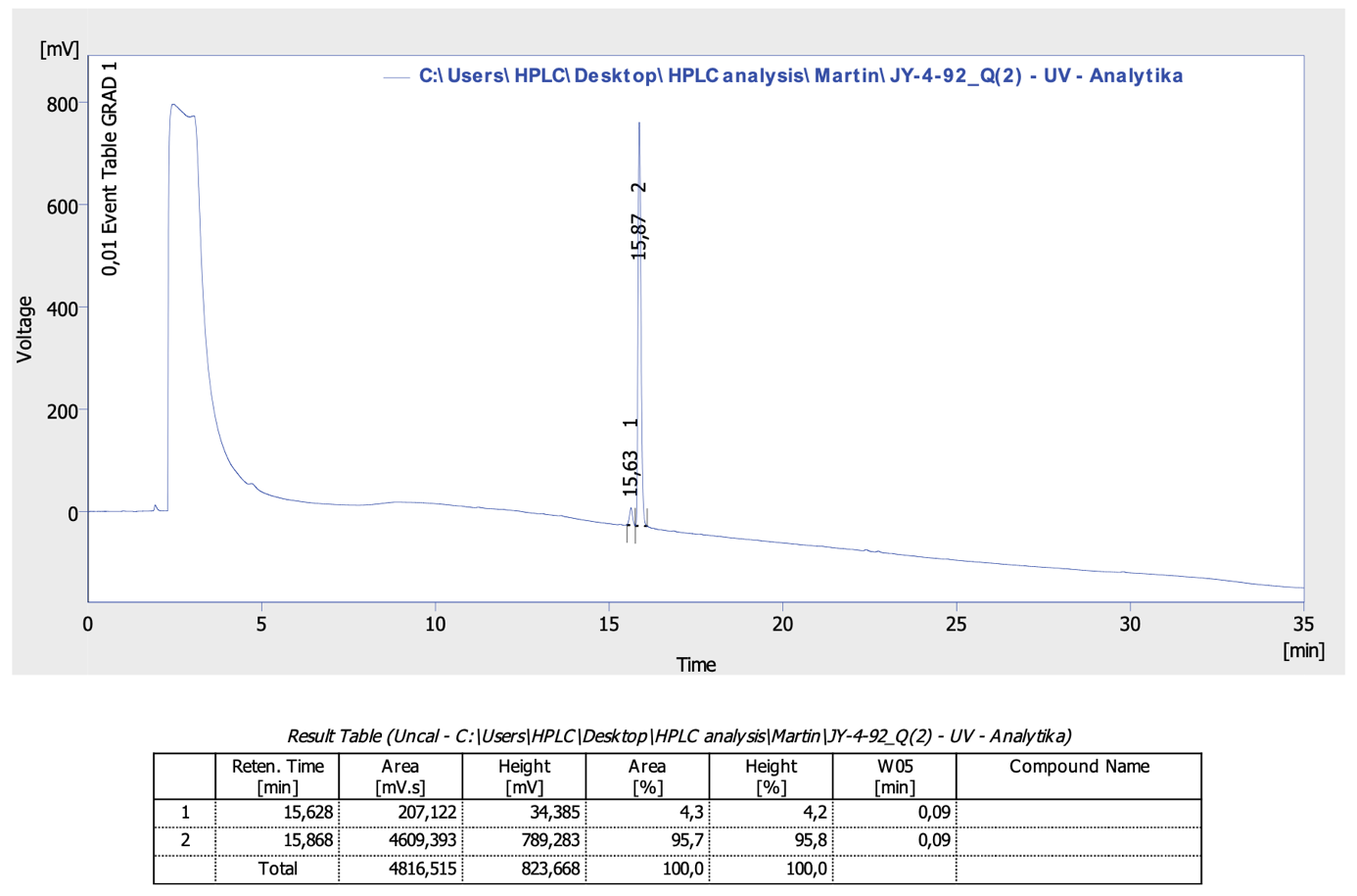

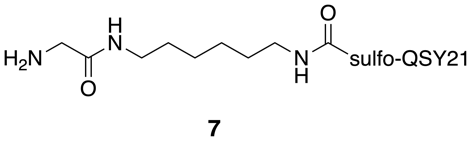

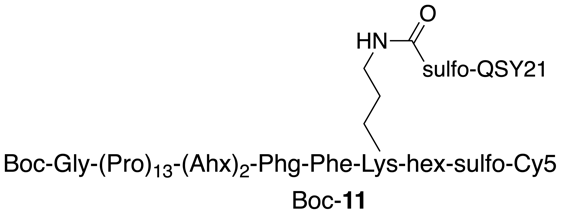

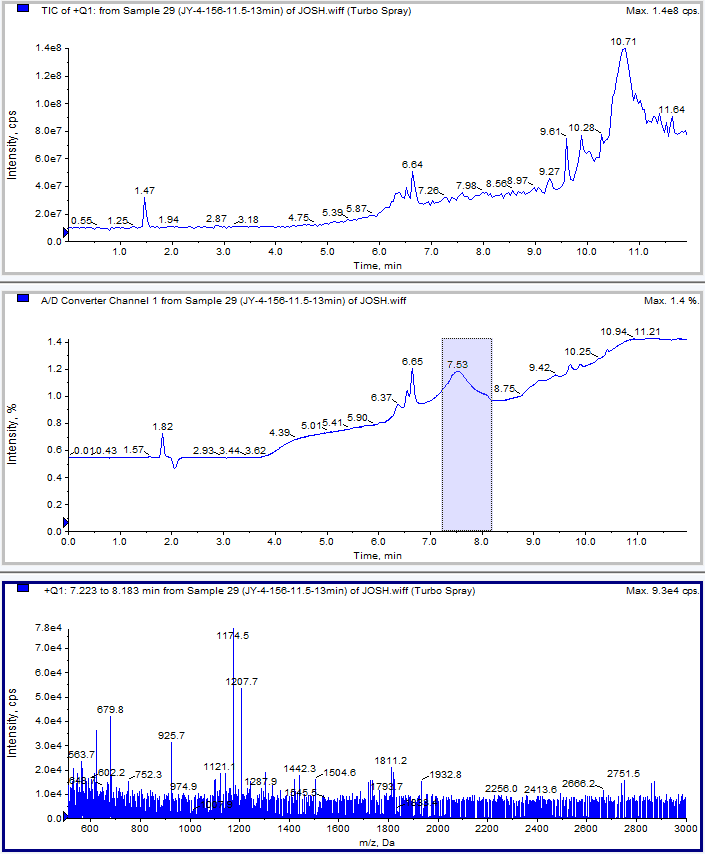

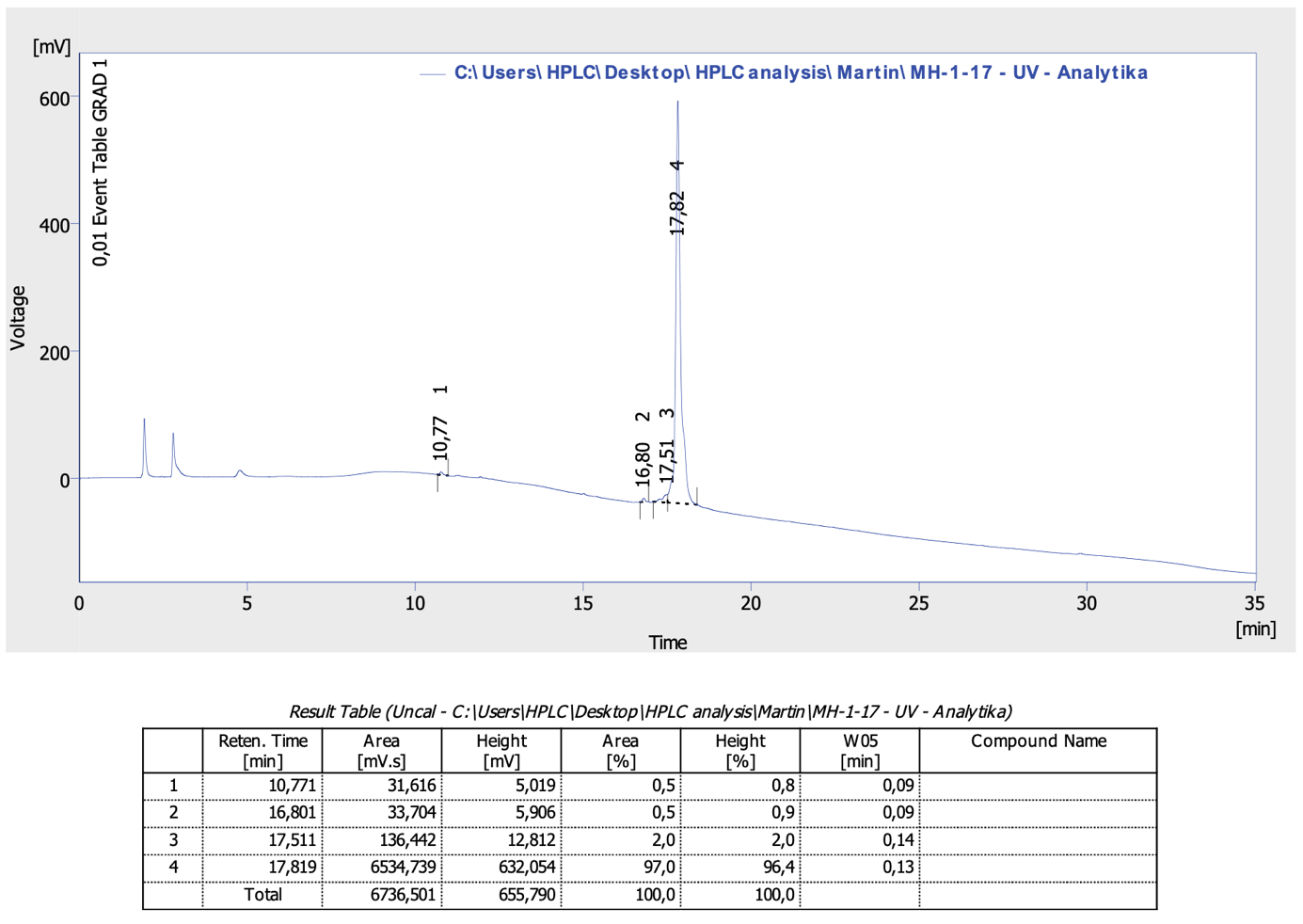

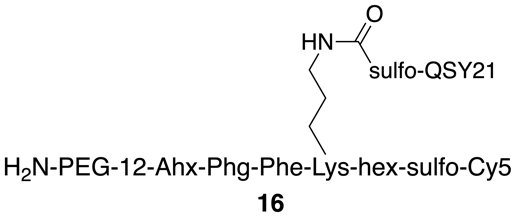

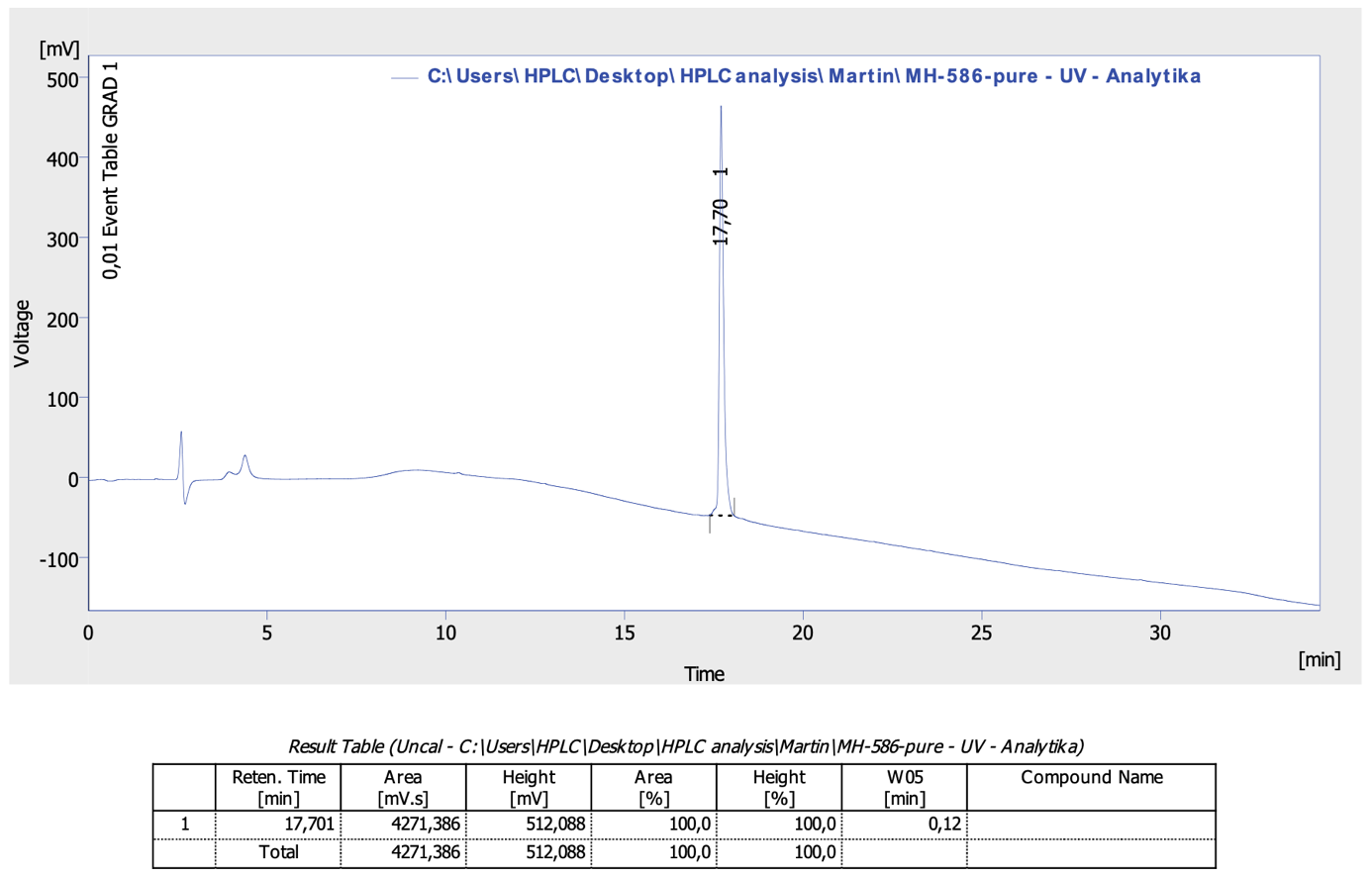

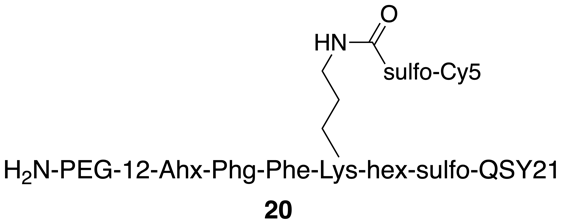

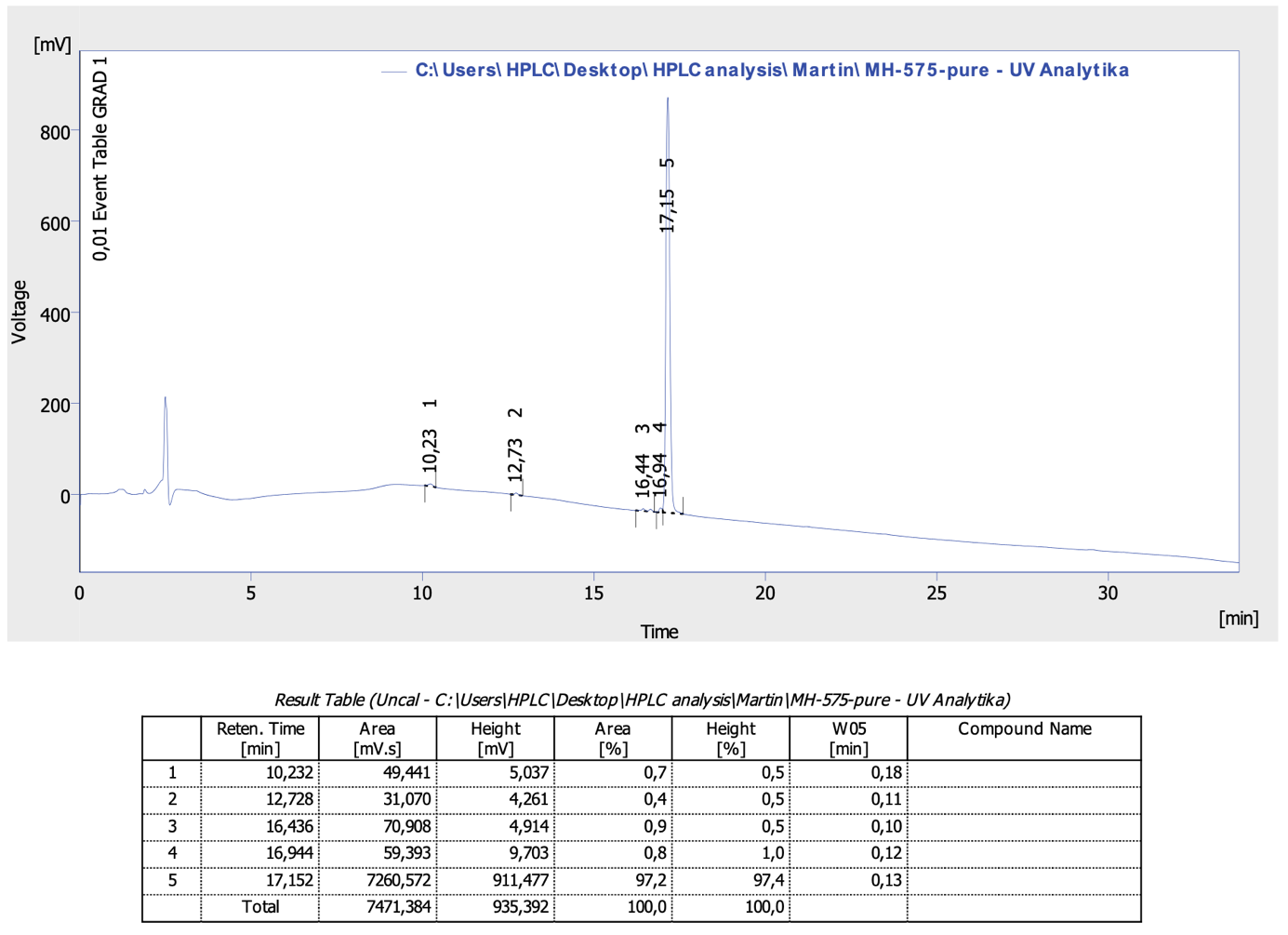

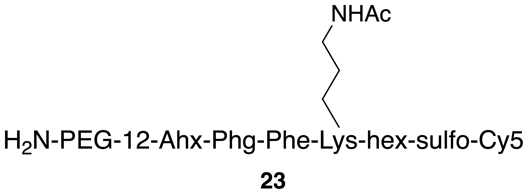

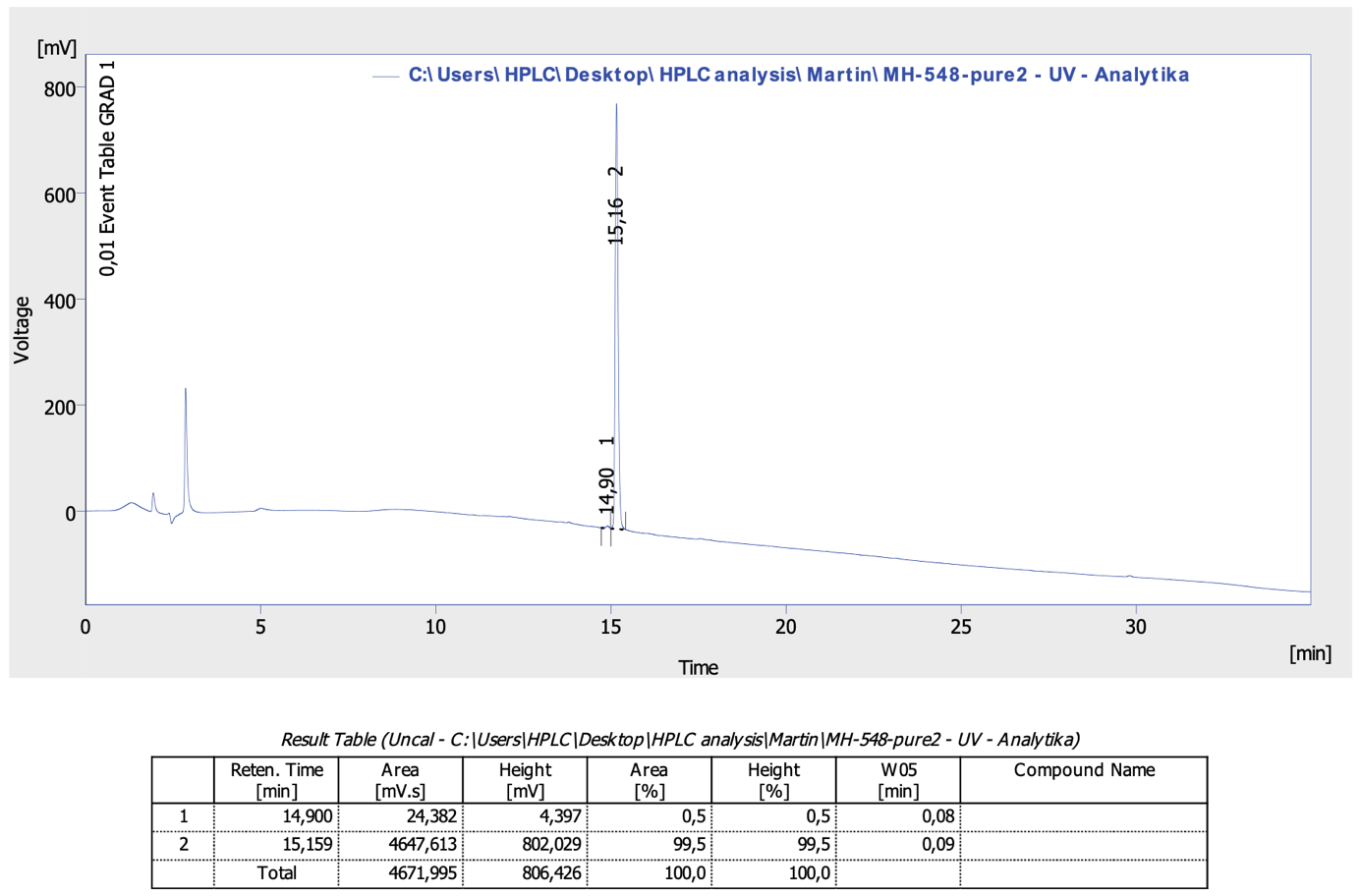

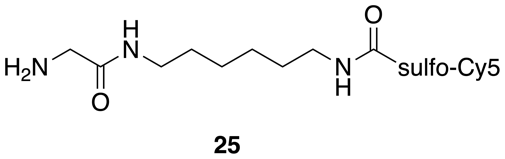

#### Synthesis of Monomers, Copolymer Precursors, and Copolymer Conjugates

**Synthesis of Monomers**

***N*-(2-hydroxypropyl)methacrylamide (HPMA):** The monomer *N*-(2-hydroxypropyl)methacrylamide (HPMA) was prepared according to a previously published method with modification.^2^ Synthetic procedure: 20.8 g (0.28 mol) of 1-amino-2-propanol (AP) was dissolved in 100 mL of ethyl acetate. The inhibitor 2,5-di-*tert*-butylhydroquinone 0.1 g was added to the solution. 42.6 g (0.28 mol) of methacryloyl anhydride was diluted with 50 mL of ethyl acetate, and 0.1 g of inhibitor was added. The methacryloyl anhydride solution was added dropwise to the AP solution at room temperature so that the temperature did not exceed 25 ℃. After adding all the methacryloyl anhydride, a homogeneous solution was formed, which was further stirred for 1 hour at laboratory temperature. After 30 min of stirring, HPMA crystals began to fall out. Then the solution was cooled to -20 ℃ with constant stirring. Product was filtered off and washed twice with diethyl ether. Finally, the monomer was recrystallized from acetone 32.5 g/70 mL acetone. Reaction yield: 29.9 g (76%).

**3-(3-methacrylamido-propanoyl)thiazolidine-2-thione (Ma-β-Ala-TT):** Monomer 3-(3-methacrylamido-propanoyl)thiazolidine-2-thione (Ma-β-Ala-TT) was prepared according to the published procedure.^3^

**Synthesis of S-2-cyano-2-propyl S'-ethyl trithiocarbonate (CTA-AIBN):** Chain transfer agent *S*-2-cyano-2-propyl *S*'-ethyl trithiocarbonate (CTA-AIBN) was prepared according to the published procedure.^4^

**Synthesis of Copolymer Precursors by RAFT Copolymerization**

Copolymer precursors pHPMA-co-Ma-β-Ala-TT (**P1–P3**) were prepared by reversible addition-fragmentation chain-transfer copolymerization (RAFT) of HPMA and Ma-β-Ala-TT in the presence of chain transfer agent (CTA-AIBN) and initiator 2,2'-azobis(4-methoxy-2,4-dimethylvaleronitrile (V-70).

Briefly, synthesis of copolymer precursor **P1**: **HPMA** (1.25 g, 8.73 mmol) was dissolved in 12.0 mL of tert-BuOH and mixed with **Ma-β-Ala-TT** (0.308 g, 1.19 mmol) dissolved in 2.1 mL of DMAA. Then was added **CTA-AIBN** (3.4 mg, 1.65x10^-2^ mmol) and V-70 (2.55mg 8.27x10^-3^ mmol). Ratio of monomer/CTA/V-70 was 600/1/0.5. Polymerization mixture was bubbled with argon for 10 min and then sealed in ampule and copolymerization was carried out at 40 °C for 24 h. Copolymer was isolated by precipitation into mixture acetone:diethyl ether 3:1, filtered off and dried in vacuum. Copolymer was dissolved in methanol and reprecipitated into acetone:diethyl ether 3:1, filtered off and dried in vacuum. The trithiocarbonate terminating groups were removed by method described by Perrier.^5^ The yield was 0.90 g of copolymer **P1**.

**Synthesis of Copolymer Conjugates**

Conjugates were prepared by aminolytic reaction of polymer precursor containing TT reactive groups with ligands (quencher or fluorophore or combination of both) in DMSO in the presence DIPEA. Reaction was carried out for 4 h at rt. Residual TT groups were aminolyzed by addition of 1-amino-2-propanol and purified on LH-20 column in methanol. Methanol was evaporated, conjugate was dissolved in distilled water, and finely purified on Sephadex PD-10 column and lyophilized.

**S-QF**: Polymer precursor **P1** (11 mg, 9.42x10^-6^ mol TT groups) was dissolved in 300 µL DMSO and compound **3** (1.7 mg, 7.6x10^-7^ mol) was added followed by addition of DIPEA (0.5 µL, 3.0x10^-6^ mol). Reaction was carried out for 4 h at room temperature. Residual TT groups were aminolyzed by addition of 1-amino-2-propanol (10 µL). reaction mixture was diluted with 1 mL of methanol and purified on LH-20 column 1.5 x 20 cm in methanol. Methanol was evaporated, conjugate was dissolved in 1.5 mL distilled water and finely purified on Sephadex PD-10 column and lyophilized. The yield of conjugate was 10.7 mg.

**S-Q1:1F:** Polymer precursor **P1** (11 mg, 9.42x10^-6^ mol TT groups) was dissolved in 300 µL DMSO and compound **5** (0.7 mg, 4.9x10^-7^ mol) and compound **7** (0.7 mg, 7.0x10^-7^ mol) was added followed by addition of DIPEA (0.8 µL, 4.7x10^-6^ mol). Reaction and purification were carried out as described above. The yield of conjugate **S-Q1:1F** was 10.2 mg.

**S-Q3:1F:** Polymer precursor **P1** (11 mg, 9.42x10^-6^ mol TT groups) was dissolved in 300 µL DMSO and compound **5** (0.7 mg, 4.9x10^-7^ mol) and compound **7** (1.25 mg, 1.3x10^-6^ mol) was added followed by addition of DIPEA (1.2 µL, 7.0x10^-6^ mol). Reaction and purification were carried out as described above. The yield of conjugate **S-Q3:1F** was 11.7 mg.

**P-QF:** Polymer precursor **P2** (9.1 mg, 7.6x10^-6^ mol TT groups) was dissolved in 300 µL DMSO and compound **11** (1.5 mg, 4.3x10^-7^ mol) was added followed by addition of DIPEA (0.7 µL, 4.3x10^-6^ mol). Reaction and purification were carried out as described above. The yield of conjugate **P-QF** was 9.9 mg.

**L-QF:** Polymer precursor **P3** (8.1 mg, 6.0x10^-6^ mol TT groups) was dissolved in 300 µL DMSO and compound **16** (1.0 mg, 3.6x10^-7^ mol) was added followed by addition of DIPEA (0.6 µL, 3.6x10^-6^ mol). Reaction and purification were carried out as described above. The yield of conjugate **L-QF** was 8.6 mg.

**L-FQ:** Polymer precursor **P3** (8.1 mg, 6.0x10^-6^ mol TT groups) was dissolved in 300 µL DMSO and compound **20** (0.95 mg, 3.4x10^-7^ mol) followed by addition of DIPEA (0.6 µL, 3.4x10^-6^ mol). Reaction and purification were carried out as described above. The yield of conjugate **L-FQ** was 8.6 mg.

**L-Q1:1F:** Polymer precursor **P3** (11 mg, 9.42x10^-6^ mol TT groups) was dissolved in 300 µL DMSO and compound **23** (0.7 mg, 3.5x10^-7^ mol) and compound **7** (0.4 mg, 3.6x10^-7^ mol) was added followed by addition of DIPEA (1.2 µL, 7.1x10^-6^ mol). Reaction and purification were carried out as described above. The yield of conjugate **L-Q1:1F** was 8.5 mg.

**L-Q2:1F:** Polymer precursor **P3** (8.0 mg, 5.94x10^-6^ mol TT groups) was dissolved in 300 µL DMSO and compound **23** (0.7 mg, 3.5x10^-7^ mol) and compound **7** (1.15 mg, 1.0x10^-6^ mol) was added followed by addition of DIPEA (2.4 µL, 1.38x10^-5^ mol). Reaction and purification were carried out as described above. The yield of conjugate **L-Q2:1F** was 8.6 mg.

**nQ:** Polymer precursor **P3** (15.1 mg, 11.2x10^-6^ mol TT groups) was dissolved in 300 µL DMSO and compound **25** (0.6 mg, 6.6x10^-7^ mol) followed by addition of DIPEA (0.6 µL, 3.4x10^-6^ mol). Reaction and purification were carried out as described above. The yield of conjugate **nQ** was 14.2 mg.

**Characterization of Monomers, Polymer Precursors and Polymer Conjugates**

Monomers and chain transfer agents were characterized on an HPLC system (Shimadzu, Japan) equipped with a UV/Vis photodiode array detector and a reverse-phase column (Chromolith High-Resolution RP-18e, 100x4.6 mm) (Merck, Czech Republic). The mobile phase was 5-95% water-acetonitrile gradient with 0.1% TFA for 15 min at a flow rate of 1.0 mL·min^-1^. The molar mass of the monomers and CTA was determined by mass spectrometry (MS LCQ Fleet, Thermo Fisher Scientific).

The content of TT reactive groups was determined by spectrophotometric analysis on a SPECORD 205 (Jena Analytics, Germany) at 305 nm (ε_305_ = 10,800 L·mol^−1^·cm^−1^; methanol); the content of the ligand with Cy5 in polymer conjugate (**nQ**) was determined at 646 nm (ε_646_ = 250,000 L·mol^−1^·cm^−1^; distilled water). Polymer precursors were characterized by number-average molecular weight (*M*_n_), weight-average molecular weight (*M*_w_), and polydispersity (*Đ*) by size exclusion chromatography (SEC) on a system consisting of HPLC (Shimadzu, Japan) equipped with a UV detector and Wyatt Technology detectors: Optilab rEX differential refractometer and DAWN 8 multiangle light scattering detector. Column TSKgel G4000SW with 20% 0.3 M acetate buffer (pH 6.5)/ 80% methanol (v/v) as a mobile phase was used. The content of the ligands in the polymer conjugates was determined in the hydrolysate (6 N HCl, 115°C, 16 h) using HPLC (Shimadzu, Japan) with a fluorescence detector (Ex. 229 nm, Em. 490 nm) on a Chromolith^TM^ RP-C18 column by the method of pre-column derivatization with
*o*-naphthalenedialdehyde.

#### 4T1 Breast Tumor Model

Prior to injecting the cells for the tumor model, 4T1 cells were cultured to confluency in treated 10 cm dishes. 4T1 cells were detached using pre-warmed Tryspin (Gibco, Cat. No.: 25300062) via incubation for 3 min at 37 °C. Cells were detached with additional pipetting after the addition of cell culture media. Cells were then centrifuged for 3 min at 250 g and the media removed. Cells were resuspended in PBS, centrifuged, and PBS removed a total of three times to wash the cells. The 4T1 cells were then resuspended in PBS to a dilution of 1 × 10^6^ cells per mL. While under isoflurane anesthesia, mice were subcutaneously injected in the third and eighth mammary fat pads of BALB/c female mice (aged 6–8 weeks; Jackson Laboratory) with 100 μL of the diluted 4T1 cells (1 × 10^5^ cells per fat pad). The seeded tumors were allowed to grow for 7-10 days. Body fur was removed using Nair lotion in the mammary fat pads and surrounding area while under anesthesia prior to probe injection. Once the tumors were developed, mice were injected intravenously with 6.25 nmols of probe in 100 μL using a 28-gauge 1 mL insulin syringe into the tail vein. **6-QC** was dissolved in a 100 μL formulation of 1× PBS with 10% DMSO and 30% PEG 400. All macromolecular probes were dissolved in a formulation of 1× PBS with 10% DMSO. Post injection, the mice were non-invasively imaged using the LI-COR Pearl Trilogy imaging system at multiple time points. The 700 nm channel and accompanying standard excitation/emission settings were used to image Cyanine5 fluorescence along with the standard white light channel filters at a resolution of 85 µm. After the live imaging, mice were euthanized using cervical dislocation under isoflurane anesthesia. Mice were then splayed and imaged. Finally, through dissection organs were collected (liver, kidney, spleen, lung, heart) as well as the primary tumors and non-injected fat pad controls. These tissues were then imaged *ex-vivo*. Fluorescence intensity was measured using the built-in LI-COR Image Studio software. To quantify the tumor to background ratio in the live animals and splayed images, the average fluorescence intensity in the tumor was divided by the average fluorescence intensity of adjacent healthy tissue using an ellipsoid region of interest of the same size. For quantification of ex-*vivo* tissues, the average fluorescence intensity was measured in the tissue of interest using the largest ellipsoid region of interest possible for each tissue. Fluorescence intensity color plots were generated using the built in LI-COR Image Studio software. All images shown were linked to display the same brightness and contrast settings for a given condition (example: 24-hour images for all probes are linked). Plots were generated and statistics evaluated using GraphPad Prism 10.2.0 (392).

### References

1.	Ofori, L. O.; Withana, N. P.; Prestwood, T. R.; Verdoes, M.; Brady, J. J.; Winslow, M. M.; Sorger, J.; Bogyo, M. Design of Protease Activated Optical Contrast Agents That Exploit a Latent Lysosomotropic Effect for Use in Fluorescence-Guided Surgery. *ACS Chem. Biol*. 2015, 10 (9), 1977–1988.

2. Fairbanks, B.D.; Thissen, H.; Maurdev, G.; Pasic, P.; White, J. F.; Meagher, L. Inhibition of Protein and Cell Attachment on Materials Generated from N-(2-Hydroxypropyl) Acrylamide, *Biomacromolecules* 2014, 15(9), 3259-3266.

3.	Šubr, V.; Ulbrich, K.; Synthesis and properties of new N-(2-hydroxypropyl)methacrylamide copolymers containing thiazolidine-2-thione reactive groups, React. *Funct. Polym*. 2006, 66, 1525-1538.

4.	Ishitake, K.; Satoh, K.; Kamigaito, M.; Okamoto, Y.; Stereogradient Polymers Formed by Controlled/Living Radical Polymerization of Bulky Methacrylate Monomers, *Angew. Chem. Int. Ed.* 2009, 48(11) (2009) 1991-1994.

5.	Perrier, S.; Takolpuckdee, P.; Mars, C.A. Reversible addition-fragmentation chain transfer polymerization: End group modification for functionalized polymers and chain transfer agent recovery, *Macromolecules* 2005, 38(6), 2033-2036.
